# Game structure shapes individuals’ norms in collective decision-making

**DOI:** 10.1101/2025.10.17.683048

**Authors:** Yang Li, Xin Wang, Wucheng Dong, Xingru Chen, Hongwei Zheng, Longzhao Liu, Shaoting Tang

## Abstract

Collective decision-making often requires individuals to reconcile private benefits with collective benefits of cooperation. Norms can sustain the cooperation by shaping how individuals evaluate consequences of actions and respond to shared behavioral standard. However, whether different game structures give rise to different norms, and the specific mechanisms underlying such effects, remain unclear. Here, we compared linear and threshold public goods games (LPGGs and TPGGs) in repeated four-person interactions, combining model-free behavioral analysis, agent-based modelling and fNIRS hyperscanning. LPGG produced more dynamic contribution updating and was more influenced by others’ recent behavior, whereas TPGG showed more stable consensus and stronger punishment enforcement. Agent-based modelling revealed a corresponding dissociation: LPGG placed greater weight on consequence-based social norms, whereas TPGG showed stronger conformity-based norms. These normative differences were accompanied by distinct neural characteristics. TPGG elicited stronger rTPJ activation, which was positively associated with contributions. dmPFC–rTPJ connectivity tracked distinct consequence-based social parameters, namely the baseline weight in TPGG and context-dependent modulation in LPGG. At the group level, interaction-specific dmPFC inter-brain neural synchrony emerged only in TPGG but not in LPGG. Together, these findings show that game structure does more than alter incentives: it reorganizes the behavioral, computational and neural mechanisms through which norms sustain collective cooperation.

## Introduction

Collective action often confronts individuals with a fundamental social dilemma: actions that maximize private benefits can undermine outcomes that are collectively beneficial. One mechanism through which groups can overcome this tension and sustain cooperation is the social norms [1–3]. Social norms can be understood as shared expectations about how group members should behave, supported by beliefs about both others’ behavior and the social appropriateness of alternative actions [1, 4, 5]. By defining which behaviors are expected or socially approved, social norms can stabilize cooperation even when private incentives favor defection [2, 3, 6].

Norms do not operate through a single route [5, 7, 8]. An important unresolved question is therefore whether the structure of collective decision-making determines which of these norms governs behavior [1, 3, 5, 9]. In some collective settings, norms may be internalized as preferences for actions that generate beneficial consequences for others [2, 7], whereas others may encourage individuals to conform to a behavioral standard [1, 4, 10]. Identifying which norm predominates under different game structures is essential not only for explaining how cooperation is maintained, but also for predicting how collective behavior will respond when environment changes. Linear and threshold public goods games (LPGGs and TPGGs) provide a particularly useful contrast for testing how collective structure reorganizes different norms as many laboratory game paradigms, including step-level public goods dilemmas [11], are primarily derived from two foundational structures [12–14]. Both games preserve the core social dilemma in which contribution is privately costly but collectively beneficial, thereby retaining the scope for free-riding [15, 16]. Yet they differ in how individual actions translate into collective outcomes. In the LPGG, every contribution continuously increases the collective return. Individuals therefore benefit from others’ contributions while retaining motives to reduce their own. By contrast, collective benefits are realized only when total group contributions reach a predefined threshold in the TPGG.

Norms emerge through repurposing of basic cognitive and neural processes [17]. It is acknowledged that social decision-making recruits regions including the right temporoparietal junction (rTPJ) and dorsomedial prefrontal cortex (dmPFC), which support the representation and integration of socially relevant information [18–20]. Hyperscanning further enables neural responses to be examined simultaneously across interacting individuals [21, 22], with inter-brain neural synchrony (INS) providing a measure of interpersonal neural alignment during collective behavior [23, 24]. These measures can test whether norms across game structures is further accompanied by corresponding differences in individual neural responses and group-level neural alignment [24, 25].

In this study, we compared the LPGG and TPGG within a unified four-person repeated public goods game paradigm while simultaneously recording neural responses using fNIRS hyperscanning [26]. The two types of experiment were matched in contribution options and group composition but differed in whether collective returns changed continuously with total contribution or depended on reaching a predefined threshold. We further introduced heterogeneous productivity [27–29] and peer punishment [30–32] to ensure that our findings generalize to real-world contexts with different work efficiency and institutional environments. To identify the latent norms, we developed a utility-based computational framework in which each candidate action was evaluated through three separable components [33–35]. Self-regarding material utility captured the consequences of an action for the focal individual’s own payoff. Consequence-based social utility captured an internalized normative concern for the material consequences that one’s action generated for other group members [2, 7, 36]. Its weight could either remain fixed or vary with the recent social context, allowing us to identify the context-dependent modulation [9, 37]. Conformity-based utility captured the subjective cost associated with deviating from an inferred standard based on the recent behavior of others [4, 10].

Across behavioral, computational, and neural levels, our findings converged on a common distinction between the LPGG and TPGG [1, 5]. Behavior in LPGG showed stronger flexibility and was more strongly influenced by others’ contributions, whereas behavior in TPGG showed stronger stickiness to individuals’ own previous contributions and was accompanied with stronger punishment enforcement. Computational model comparison revealed a corresponding shift in dominant norm. LPGG was best captured by *S*-*vCS*-*C* model, and consequence-based social utility was higher than in TPGG. In contrast, TPGG was best captured by *S*-*f CS*-*C* model, while conformity-based utility exerted a stronger influence on contribution decisions. Neural measures further reinforced this distinction. TPGG elicited stronger rTPJ activation, which was positively associated with contributions. Functional connectivity between rTPJ and dmPFC tracked distinct consequence-based social components, namely the baseline social weight in TPGG and context-dependent modulation in LPGG. At the group level, significant dmPFC INS emerged in TPGG but not LPGG, further indicating that the TPGG was consistent with stronger neural alignment. Together, these results suggest that game structure does more than alter incentives: it reorganizes the latent norms [5]. LPGG favours flexible, consequence-based social values, whereas TPGG stabilizes conformity and is accompanied by stronger neural alignment for collective decision-making [12, 38].

## Results

### Linear and threshold public goods games

To formalize the two game structures, let 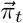 denote the vector of individual payoffs in round *t*, 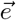 the vector of individual endowments, 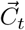 the vector of individual contributions, and ***P*** the diagonal productivity matrix. Here, 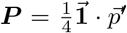 with 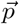 the productivity vector (Figure 1, a, b). The payoff vector was defined as:

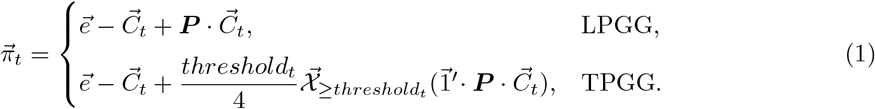

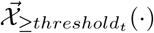 denotes an indicator vector that takes value 1 when the threshold condition is satisfied and 0 otherwise. Thus, in LPGG, weighted contributions were transformed continuously into returns, whereas in TPGG, returns were generated only when the group’s total weighted contributions, 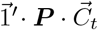, reached the threshold *threshold*_*t*_.

**Fig. 1:**
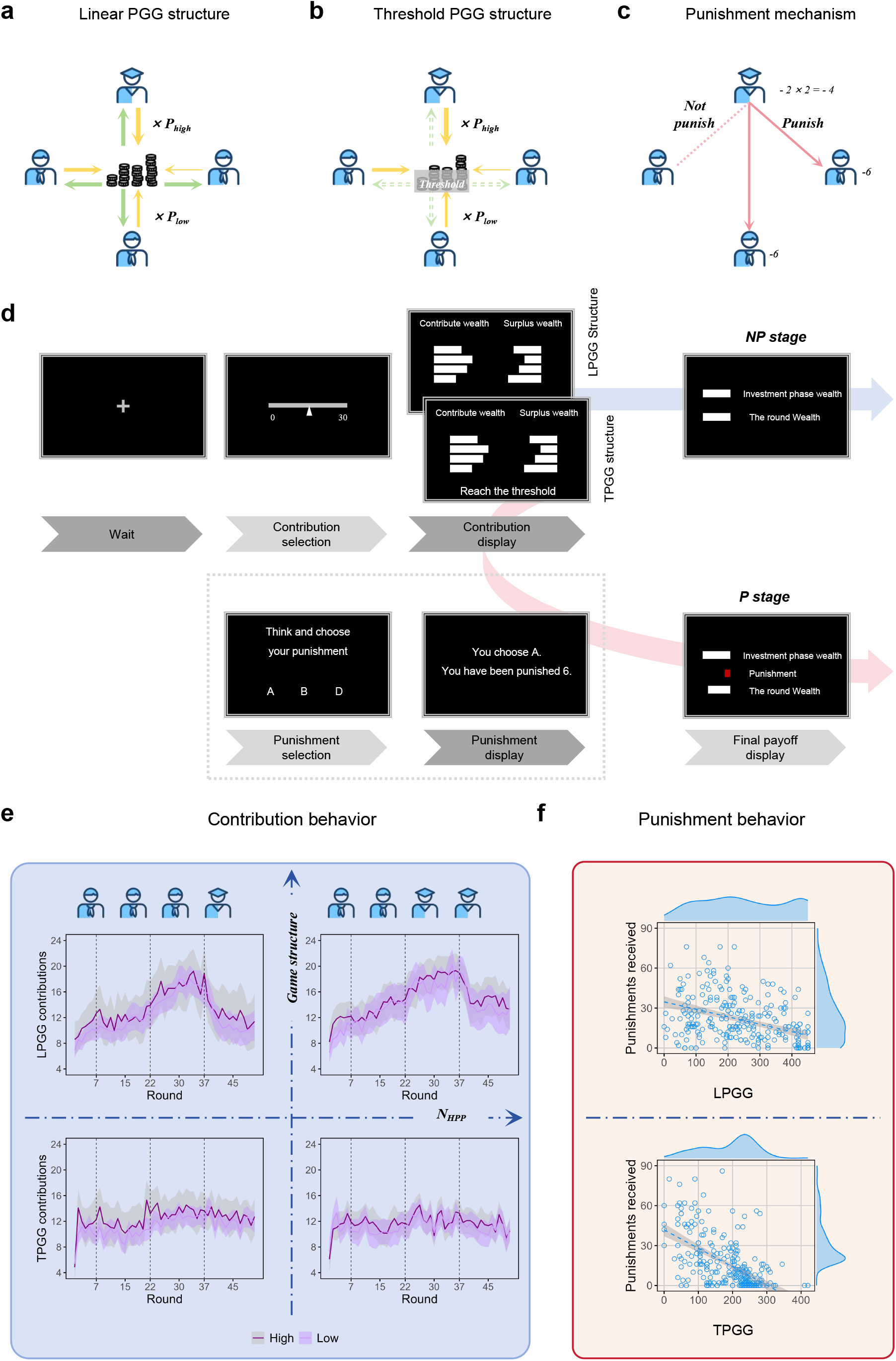
Game structures, experimental procedure and model-free behavior. **a,b**, Schematic representations of the linear public-goods game (LPGG) and threshold public-goods game (TPGG). In both games, four participants contributed to a common pool under heterogeneous productivity; collective returns increased continuously with productivity-weighted contributions in LPGG (**a**), whereas returns in TPGG were realized only when the group reached the threshold (**b**). **c**, Peer punishment. During punishment phases, participants could assign punishment to other group members after observing their contributions. **d**, Trial structure. Each trial comprised a fixation period, contribution selection and contribution outcome. During P stages, these phases were followed by punishment selection and punishment outcome before the final payoff was presented; during NP stages (NP), the punishment phases were omitted. **e**, Round-by-round contribution dynamics in LPGG and TPGG. Lines denote mean contributions of high-productivity participants (HPPs) and low-productivity participants (LPPs), and shaded regions indicate 95% confidence intervals. Vertical dashed lines mark transitions between the preparation, NP1, P and NP2 stages. **f**, Relationship between total contributions and punishments received during the P stage. Each point represents one participant; fitted lines show linear regressions with 95% confidence intervals, and marginal distributions show the corresponding distributions of total contributions and punishments received.

Punishment was implemented after contribution (Figure 1, c). Individual *i* could impose punishment points (i.e., punishments) at a cost of 2 points to reduce the payoff of any other group member by 6 points

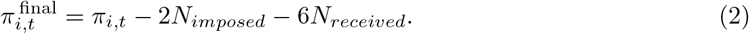

Here, *N*_*imposed*_ and *N*_*received*_ denote the number of individuals punished by individual *i* and the number of individuals who punished individual *i*, respectively.

We employed a 2 *×* 2 mixed factorial design: between-subjects factor (LPGG vs. TPGG) and within-subjects factor ([peer] punishment vs. no [peer] punishment). Further, we manipulated the number of high-productivity participants (*p* = 3.2) across treatments (*n*_*HPP*_ = 1, 2). A total of 540 participants were randomly assigned to 135 four-person groups and completed a real-time interactive task. During the experiment, all participants wore multi-channel fNIRS headsets to record hemo-dynamics from predefined brain regions of interest. The task was an repeated PGG consisting of three phases per round: investment, punishment (P stage), and outcome display (Figure 1, d). Each participant received an endowment of 30 points at the beginning of each round and simultaneously selected one of seven possible contributions on the slider (0, 5, 10, 15, 20, 25, or 30 points). Each group completed 51 rounds divided into four stages: preparation stage (rounds 1-6), first no punishment stage (NP1, rounds 7-21), punishment stage (P, rounds 22-36), and second no punishment stage (NP2, rounds 37-51). The introduction of NP2 stage ruled out order effects, confirming that peer punishment, not round order, affected the contributions [30]. The entire experimental procedure was conducted on the day of the main experiment and lasted approximately 100 minutes. This duration encompassed pre-experiment assessment and instructions, main experiment, and post-experiment survey (see Method Main experiment for full details).

### Game structures shape different contribution dynamics

We first presented the mean contributions with 95% confidence intervals in each round of high-productivity and low-productivity participants (Figure 1, e). No round-by-round differences of contributions between high- and low-productivity individuals survived FDR correction in either LPGG or TPGG. Moreover, the two productivity individuals exhibited similar behavioral dynamics across rounds. On this basis, we pooled the two productivity individuals in subsequent analyses. We next examined how contributions evolved across rounds. We modeled contributions using a linear mixed-effects model, with round, peer punishment availability, and their interaction entered as fixed effects. Participant identity was included as a random intercept.

In LPGG, punishment increased contributions (LPGG 1H: β = 4.625, SE = 0.203, *t*_5629_ = 22.831, *P <* 0.001; LPGG 2H: β = 3.733, SE = 0.215, *t*_5629_ = 17.364, *P <* 0.001) and showed a significant punishment-by-round interaction (LPGG 1H: β = 0.418, SE = 0.047, *t*_5629_ = 8.920, *P <* 0.001; LPGG 2H: β = 0.234, SE = 0.050, *t*_5629_ = 4.711, *P <* 0.001). By contrast, in TPGG, punishment also increased contributions (TPGG 1H: β = 1.247, SE = 0.181, *t*_5805_ = 6.900, *P <* 0.001; TPGG 2H: β = 1.103, SE = 0.186, *t*_5629_ = 5.934, *P <* 0.001), but its impact on behavioral dynamics was weaker (TPGG 1H: β = 0.118, SE = 0.042, *t*_5805_ = 2.821, *P* = 0.0048; TPGG 2H: β = 0.055, SE = 0.043, *t*_5629_ = 1.283, *P* = 0.200). Simple-slope analyses during P stage showed significant positive trajectories in LPGG (LPGG 1H: β = 0.306, SE = 0.038, *t*_5629_ = 7.984, 95% CI [0.231, 0.381], *P <* 0.001; LPGG 2H: β = 0.229, SE = 0.041, *t*_5629_ = 5.638, 95% CI [0.149, 0.309], *P <* 0.001).

In TPGG, the trajectory was positive in TPGG 1H (β = 0.087, SE = 0.034, *t*_5805_ = 2.542, 95% CI [0.020, 0.154], *P* = 0.011), but not in TPGG 2H (β = *−*0.005, SE = 0.035, *t*_5629_ = *−*0.143, 95% CI [*−*0.074, 0.064], *P* = 0.886). We next examined aggregate punishment. The total punishment an individual received decreased with their total contribution (Figure 1, f), consistent with greater punishment of lower contributors. This negative association was evident in both game structures (LPGG: β = *−*0.056, SE = 0.007, *t*_254_ = *−*7.688, 95% CI [*−*0.070, *−*0.042], *P <* 0.001; TPGG: β = *−*0.138, SE = 0.012, t_258_ = *−*11.530, 95% CI [*−*0.162, *−*0.115], *P <* 0.001). Furthermore, the estimated slope was larger in magnitude in TPGG than in LPGG.

Conditional contribution patterns further revealed distinct modes of behavioral dynamics across game structures [14, 39, 40] (Figure 2, see Methods Analyses of conditional behavior). Contributions in both games depended positively on individuals’ own previous contributions (*stickiness*) and those of other group members (*imitation*), with punishment increasing stickiness in both structures. Critically, TPGG consistently showed greater stickiness (NP: β = 0.38, 95% CI [0.34, 0.42]; P: β = 0.42, 95% CI [0.36, 0.47]) but weaker imitation (NP: β = 0.35, 95% CI [0.31, 0.38]; P: β = 0.34, 95% CI [0.29, 0.38]) than LPGG (stickiness, NP: β = 0.30, 95% CI [0.26, 0.34]; P: β = 0.38, 95% CI [0.33, 0.44]; imitation, NP: β = 0.43, 95% CI [0.39, 0.46]; P: β = 0.39, 95% CI [0.35, 0.44]). Stickiness and imitation were also negatively associated across individuals (Supplementary Figure 7), and the pattern remained unchanged when contributions were multiplied by the corresponding productivity (Supplementary Figure 8). Overall, these model-free analyses revealed distinct behavioral organization across game structures: LPGG showed higher flexibility and was more influenced by others, whereas TPGG was characterized by more stable consensus and stickiness to own previous behavior. These behavioral signatures motivated our subsequent computational analysis of the latent norms.

**Fig. 2:**
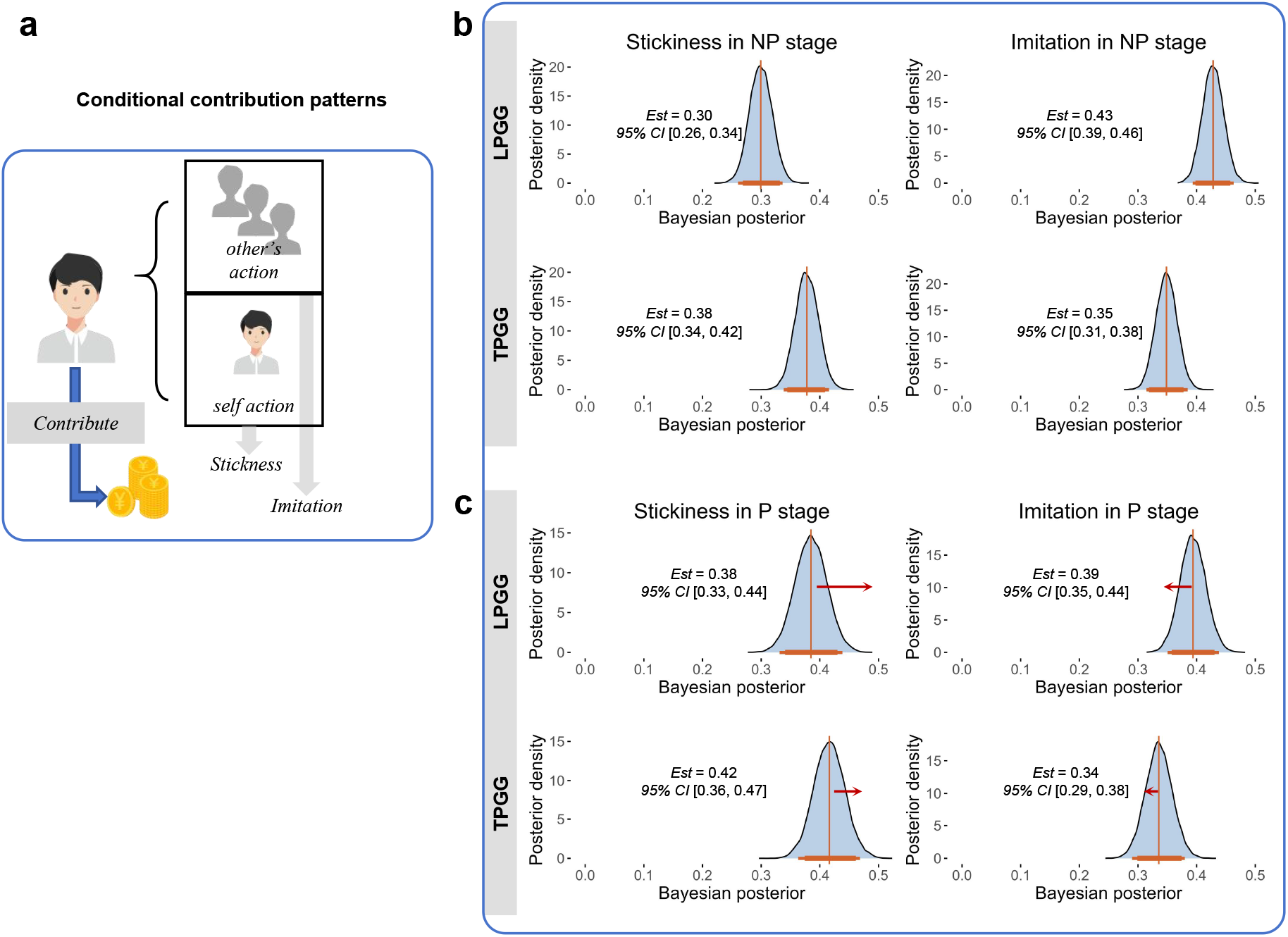
Conditional contribution patterns across game structures. **a**, Schematic of conditional contribution patterns. Contributions in the current round were modelled as a function of the focal participant’s own contributions and the contributions of others in the preceding round. The corresponding regression coefficients were interpreted as *stickiness* and *imitation*. **b**, Posterior distributions of stickiness and imitation coefficients in NP stages. TPGG showed greater stickiness but weaker imitation than LPGG. Curves show posterior distributions, with posterior estimates and 95% credible intervals indicated. **c**, Posterior distributions of stickiness and imitation coefficients in the P stage. Punishment availability increased stickiness in both game structures, while imitation remained higher in LPGG and stickiness remained higher in TPGG.

### Computational modelling revealed distinct norms across game structures

The behavioral analyses above showed that contribution dynamics differed substantially between the two game structures. These behavioral dynamics, however, do not by themselves reveal the latent norms underlying decisions. We therefore developed agent-based models to determine whether the same behavior was supported by different norms across LPGG and TPGG.

We adopted Elster’s minimalist notions [41] and decomposed the subjective utility of each candidate contribution into three components (Figure 3, a). The first component, *self-regarding material utility* (*S*), represented the material consequences of a candidate action for the focal individual’s own payoff. The second component, *consequence-based social utility* (*CS*), represented the value assigned to the consequences that one’s action created for other group members [5]. We interpreted this component as an internalized concern for how one’s behavior may affect others [2]. Its weight was allowed either to remain fixed or to vary with the recent context—the contributions of others [37]. The third component, *conformity-based utility* (*C*), represented the subjective cost of behavior deviating from a socially reference standard (see Method Agent-based modelling).

**Fig. 3:**
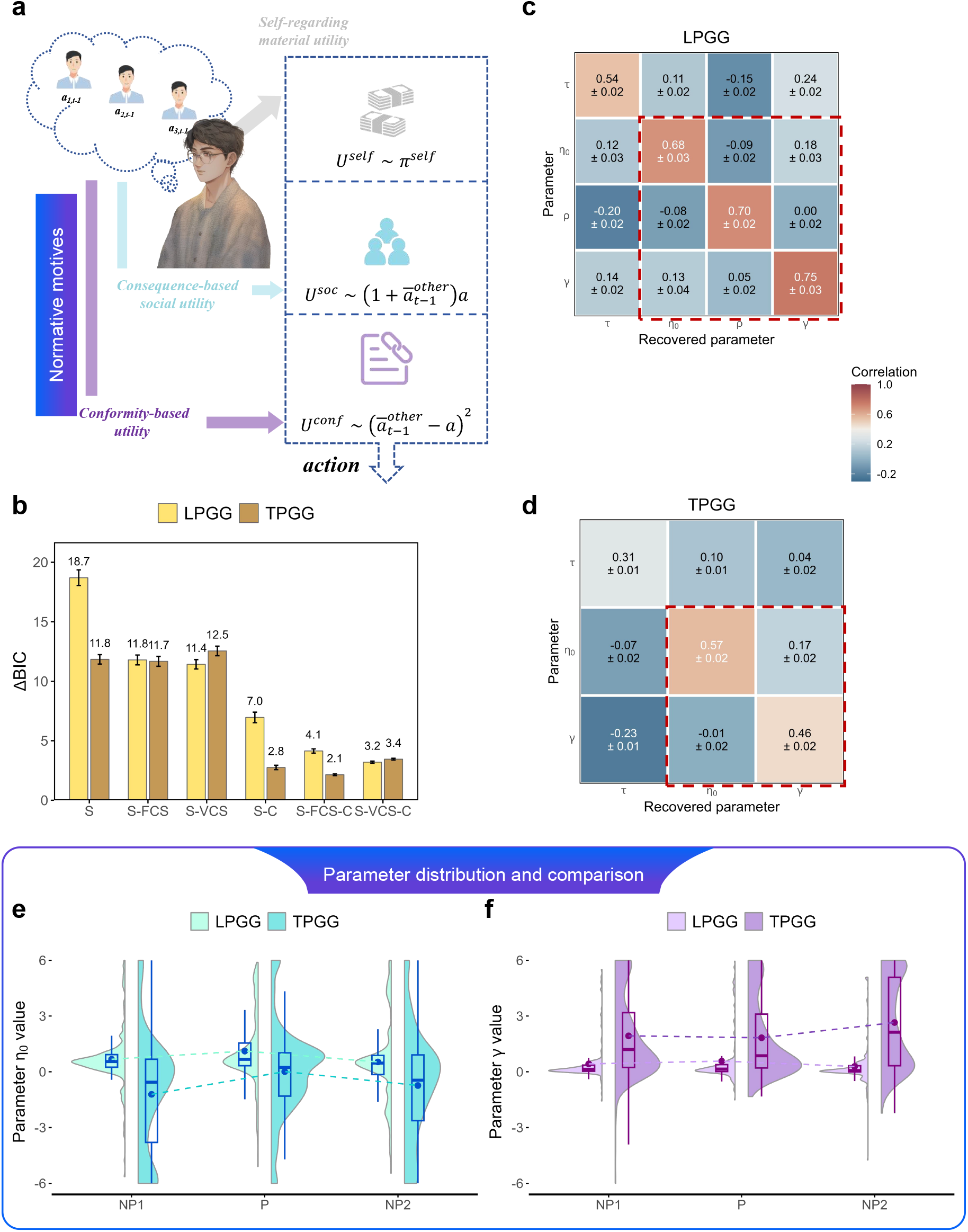
Agent-based modelling and results. **a**, Schematic of the utility-based modelling frame-work. Each candidate contribution decision was evaluated through self-regarding material utility, consequence-based social utility and conformity-based utility. **b**, Model comparison across the six candidate models. Bars show ΔBIC for LPGG and TPGG, with lower values indicating better fit. **c,d**, Parameter-recovery results for the winning LPGG (**c**) and TPGG (**d**) models. Matrix entries show the correspondence between generating and recovered parameters; diagonal elements quantify recovery of each parameter. **e**, Distribution of the baseline consequence-based social weight η_0_ across NP1, P and NP2. **f**, Distribution of the conformity-based weight γ across stages. Violin plots show parameter distributions; box plots summarize the median and interquartile range, and points indicate group means.

Our computational models were fitted to contribution behavior, while punishment availability was treated as an external environmental manipulation. Parameters were estimated separately for the NP1, P and NP2 stages. We compared six candidate models that systematically varied the presence of the *CS* and *C* components and whether the weight of *CS* was fixed or context dependent: *S, S*-*f CS, S*-*vCS, S*-*C, S*-*f CS*-*C*, and *S*-*vCS*-*C* (see Methods Utility-based models). This model space therefore allowed us to ask two distinct questions: whether contribution decisions incorporated consequence-based social motives or conformity motives beyond self-regarding material utility, and whether consequence-based social utility needed to dynamically track the recent behavior of others. For each session we computed Δ*BIC* relative to the minimum BIC among all candidate models (six in each game structure). Model comparison revealed a clear dissociation between the LPGG and TPGG (Figure 3, b). The best-fitting models were the *S*-*vCS*-*C* model for LPGG and the *S*-*f CS*-*C* model for TPGG, indicating that contribution decisions in both game structures were best explained by jointly incorporating consequence-based social and conformity-based utilities. Critically, however, the weight on consequence-based social norm required context-dependent modulation in LPGG, whereas it remained fixed in TPGG. Parameter-recovery analyses showed that simulated and recovered parameters were positively correlated along the diagonal of the recovery matrices in both structures (Figure 3, c, d). The true-recovered parameter associations were generally stronger than the off-diagonal associations, indicating that the principal motivational parameters captured distinguishable features of behavior.

The fitted parameters further revealed that LPGG and TPGG differed in the relative weight of the two norms. Across NP1, P and NP2, the baseline consequence-based social weight η_0_ was consistently higher in LPGG than in TPGG (Figure 3, e; NP1: LPGG *M* = 0.676, TPGG *M* = *−*1.21, *t*_361_ = 8.67, 95% CI [1.46, 2.32], *P <* 0.001; P: LPGG *M* = 1.12, TPGG *M* = 0.010, *t*_483_ = 5.75, 95% CI [0.728, 1.48], *P <* 0.001; NP2: LPGG *M* = 0.532, TPGG *M* = *−*0.737, *t*_477_ = 5.63, 95% CI [0.826, 1.71], *P <* 0.001). Thus, individuals assigned greater value to the consequences of their own contributions for others in LPGG. Together with the selection of the *S*-*vCS*-*C* model, this result indicates that LPGG contribution was supported by a comparatively strong consequence-based social norms whose behavioral influence remained sensitive to the recent cooperative context. The opposite pattern was observed for conformity-based weight. The γ was consistently larger in TPGG than in LPGG across all stages (Figure 3, f; NP1: LPGG *M* = 0.426, TPGG *M* = 1.93, *t*_365_ = *−*9.98, 95% CI [*−*1.80, *−*1.21], *P <* 0.001; P: LPGG *M* = 0.577, TPGG *M* = 1.83, *t*_449_ = *−*8.24, 95% CI [*−*1.55, *−*0.956], *P <* 0.001; NP2: LPGG *M* = 0.288, TPGG *M* = 2.65, *t*_360_ = *−*14.8, 95% CI [*−*2.68, *−*2.05], *P <* 0.001), which indicates that deviating from the standard implied by recent group behavior carried a larger subjective cost in TPGG. The persistence of this difference across NP1, P and NP2 further suggests that the enhanced consequence-based social norms in LPGG and conformity-based norms observed in TPGG was not merely induced by the availability of peer punishment, but reflected a more general feature of the two game structure.

### Dynamic utility trajectories revealed flexible consequence-based social utility in LPGG but conformity in TPGG

We reconstructed the trial-by-trial utility generated by the consequence-based social (*CS*) and conformity-based (*C*) components of the winning models (Figure 4, a). The two game structures exhibited distinct temporal profiles (Figure 4, b). In LPGG, *CS* utility remained positive and rose markedly when punishment became available, followed by a partial decline after punishment was removed, whereas *C* utility remained comparatively small. In TPGG, by contrast, *CS* utility remained relatively weak, whereas *C* utility showed a consistently larger negative magnitude across stages. These temporal profiles complement the parameter-level results, revealing that as interactions unfolded, behavior in LPGG became increasingly dominated by consequence-based social norms, whereas behavior in TPGG was increasingly governed by conformity-based norms.

**Fig. 4:**
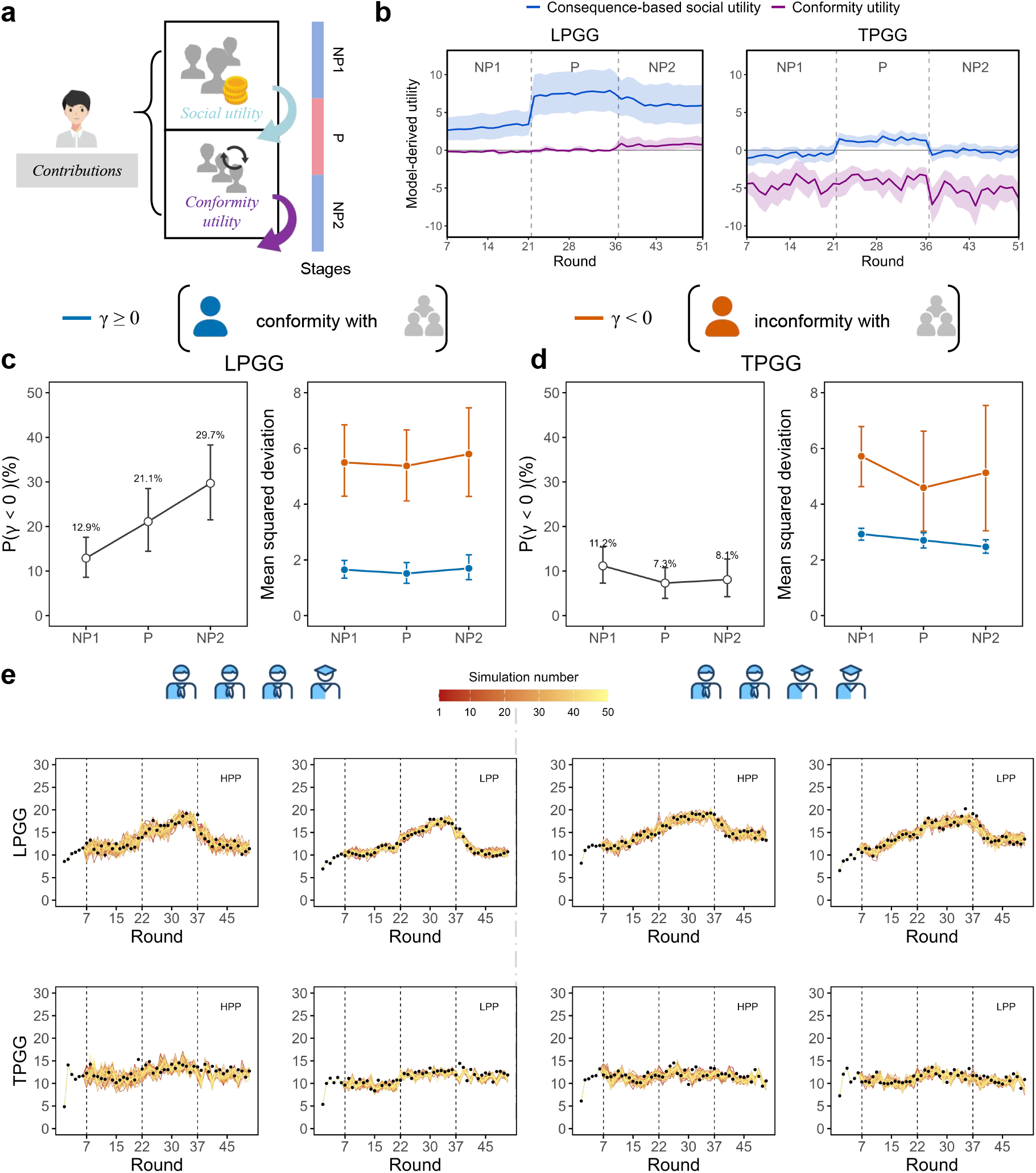
Dynamics of normative utilities, individual conformity profiles and simulations of contributions. **a**, Schematic of the trial-wise decomposition of contributions into consequence-based social (*CS*) utility and conformity-based (*C*) utility across NP1, P and NP2 stages. **b**, Round-by-round trajectories of model-derived *CS* and *C* utilities for LPGG and TPGG. Lines denote mean utility and shaded regions indicate 95% confidence intervals. **c,d**, Individual heterogeneity in conformity in LPGG (**c**) and TPGG (**d**). Participants were classified according to the sign of fitted γ. Left, proportion with γ *≥* 0 and γ < 0 across stages. Right, mean squared deviation from the standard for γ *≥* 0 and γ < 0 groups. Points indicate means and error bars denote 95% confidence intervals. **e**, Forward simulations of contribution dynamics using fitted parameters of the winning models. Coloured trajectories show 50 simulation runs, and black trajectories show the observed experimental contributions.

To understand why the average *C* utility in LPGG was slightly positive, we examined whether this pattern reflected heterogeneity in the direction of individuals’ conformity-based preferences. We examined whether the stronger conformity-based motive characterized all individuals by separating participants according to the sign of the weight γ, with γ *≥* 0 denoting the conformity with others and γ < 0 denoting the inconformity with others (Figure 4, c, d). In LPGG, the proportion of individuals with γ < 0 increased from 12.9% (95% CI [8.6%, 17.6%]) in NP1 to 21.1% ([14.5%, 28.5%]) in P and 29.7% ([21.5%, 38.3%]) in NP2, whereas it remained low and comparatively stable in TPGG (11.2% [7.3%, 15.4%], 7.3% [3.9%, 10.8%] and 8.1% [4.2%, 12.7%], respectively). Across all stages, individuals with γ < 0 also showed larger deviations from the group reference than those with γ *≥* 0 (LPGG: NP1, *M* = 5.50 vs. 1.65; P, 5.38 vs. 1.51; NP2, 5.80 vs. 1.70; TPGG: NP1, 5.72 vs. 2.93; P, 4.59 vs. 2.70; NP2, 5.13 vs. 2.47; 95% CIs shown in Figure 4, c, d). Together, these descriptive patterns suggest increasing heterogeneity in conformity-based norms over time in LPGG, whereas TPGG remained dominated by individuals with conformity. Finally, forward simulations using individually fitted parameters reproduced the distinct contribution trajectories of both productivity classes, with the *S*-*vCS*-*C* model capturing the dynamic updating in LPGG and the *S*-*f CS*-*C* model capturing the stable contributions in TPGG (Figure 4, e).

### Game structures elicited different single-brain neural characteristics and the relationship with norms

Although the social modelling results suggested that different game structures gave rise to distinct norms, neural evidence was still required to support these differences. We analyzed the time series of oxyhemoglobin concentration changes recorded by fNIRS throughout the experiment. We first compared patterns of individual neural activation between LPGG and TPGG (Figure 5, a). Significant differences emerged in channels over the right temporoparietal junction (rTPJ). Specifically, mean activation was higher in TPGG than in LPGG across four rTPJ channels (Figure 5, b; channel 5: *F*_1,506_ = 48.66, 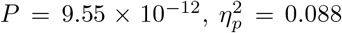, 95% CI [0.048, 0.135]; channel 6: *F*_1,496_ = 78.70, 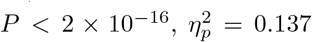, 95% CI [0.091, 0.189]; channel 7: *F*_1,505_ = 20.95, *P* = 5.93 *×* 10^*−*6^, 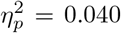, 95% CI [0.013, 0.077]; channel 8: *F*_1,503_ = 53.11, *P* = 1.23 *×* 10^*−*12^, 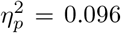, 95% CI [0.051, 0.147]; all effects survived FDR correction). Focusing on the three channels showing the strongest effects (channels 5, 6 and 8), we next examined activation during the contribution phase across the NP and P stages (Figure 5, c). Across NP1, P and NP2, mean activation in these channels remained consistently higher in TPGG than in LPGG. These findings show the greater engagement of rTPJ-related social-cognitive processes during collective decision-making in TPGG.

**Fig. 5:**
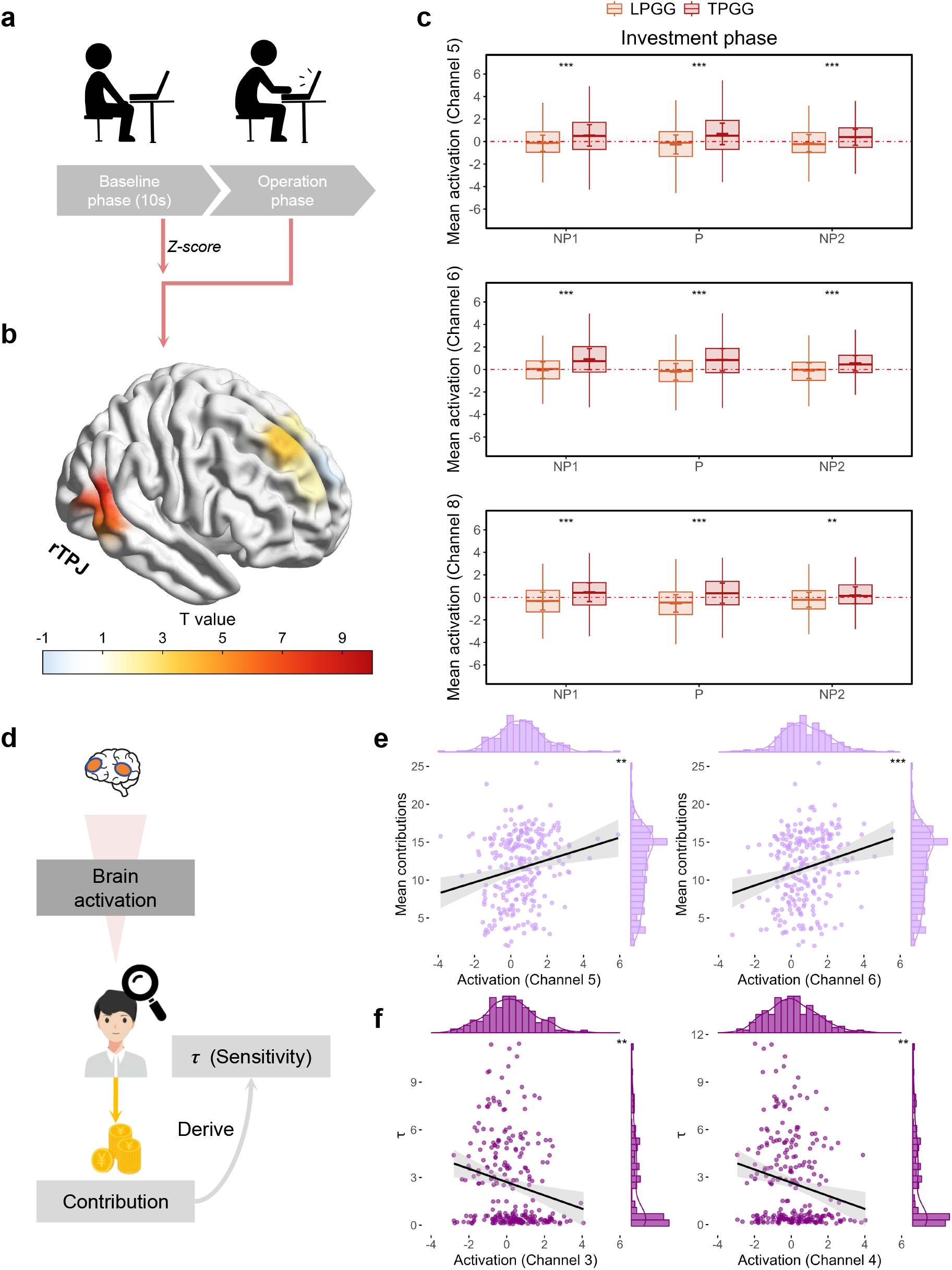
Neural activation and its relationship with behavior. **a**, Quantification of task-related neural activation. The initial 10-s interval served as the baseline, and HbO signals during the subsequent operation phases were standardized using the baseline mean and standard deviation. **b**, Differences in neural activation between LPGG and TPGG. **c**, Activation in rTPJ channels 5, 6 and 8 during contribution phases. **d**, Schematic of the individual-level analyses linking neural activation to contributions and to the inverse-temperature parameter τ. **e**, In TPGG, activation in rTPJ channels 5 and 6 was positively associated with mean contributions. **f**, In TPGG, activation in dmPFC channels 3 and 4 was negatively associated with τ. Lines denote linear fits with 95% confidence intervals; marginal distributions show the corresponding variable distributions.

To link individual neural activation with both behavior and latent decision motives, we related channel-wise fNIRS activation to two complementary individual-level measures (Figure 5, d). In TPGG, mean activation in both channel 5 and channel 6 was positively associated with mean contributions (Figure 5, e; channel 5: Pearson’s *r* = 0.206, 95% CI [0.083, 0.323], *P* = 0.0012; channel 6: *r* = 0.225, 95% CI [0.102, 0.341], *P* = 3.94 *×* 10^*−*4^), whereas no corresponding association was detected in LPGG. Thus, individuals showing stronger rTPJ engagement tended to contribute more in TPGG. We next asked whether individual neural activation was associated with the computational processes captured by the agent-based models above. We focused on the inverse-temperature parameter τ, which determines how strongly contribution decisions are governed by differences in model-derived utility: larger τ corresponds to more deterministic utility-guided decisions, whereas smaller τ reflects a flatter decision distribution. In TPGG, activation in dmPFC channels 3 and 4 was negatively associated with τ (Figure 5, e; channel 3: Pearson’s *r* = *−*0.194, 95% CI [*−*0.310, *−*0.071], *P* = 0.0021; channel 4: *r* = *−*0.196, 95% CI [*−*0.312, *−*0.074], *P* = 0.0018), whereas no corresponding association was observed in LPGG. We suggest that, in TPGG, dmPFC was recruited more strongly when individuals were less certain about which action to choose.

We next quantified within-brain functional connectivity between dmPFC (channels 1–4) and rTPJ (channels 5–8) during decision-making and related the connectivity to the fitted normative parameters (Figure 6, a–e). In TPGG, dmPFC–rTPJ connectivity was positively associated with the baseline consequence-based social weight η_0_ in TPGG (Figure 6, a,b; Ch7–Ch1: β = 0.0099, SE = 0.0032, *t*_250_ = 3.11, 95% CI [0.0036, 0.0162], *P* = 0.0021; Ch7–Ch2: β = 0.0072, SE = 0.0031, *t*_250_ = 2.31, 95% CI [0.0010, 0.0133], *P* = 0.0219; Ch5–Ch1: β = 0.0069, SE = 0.0031, *t*_250_ = 2.23, 95% CI [0.0008, 0.0130], *P* = 0.0266; Ch5–Ch2: β = 0.0074, SE = 0.0030, *t*_250_ = 2.47, 95% CI [0.0015, 0.0133], *P* = 0.0140.), whereas no corresponding associations were detected in LPGG. By contrast, in LPGG, the context-dependent modulation ρ was negatively associated with a distinct subset of dmPFC–rTPJ connections (Figure 6, d, e; Ch8–Ch1: β = *−*0.0064, SE = 0.0032, *t*_247_ = *−*1.99, 95% CI [*−*0.0126, *−*0.0001], *P* = 0.0472; Ch7–Ch4: β = *−*0.0103, SE = 0.0035, *t*_249_ = *−*2.91, 95% CI [*−*0.0173, *−*0.0033], *P* = 0.0039; Ch8–Ch4: β = *−*0.0128, SE = 0.0035, *t*_247_ = *−*3.66, 95% CI [*−*0.0196, *−*0.0059], *P* = 3.10 *×* 10^*−*4^). This counter-intuitive association may reflect a more deliberative, strategically self-regarding mode of decision-making. In LPGG, where individual payoff maximization favors lower contribution, such deliberation may shift behavior toward calculated self-interest rather than intuitive contribution, consistent with the “spontaneous giving and calculated greed” account [42]. Thus, the same within-brain functional connectivity was linked to different fitted parameters across game structures: connectivity tracked the stable baseline consequence-based social weight in TPGG, but the context-dependent modulation in LPGG. These associations suggest that game structure reorganized how social-cognitive network related to normative computation, rather than simply changing the overall strength of regional engagement.

**Fig. 6:**
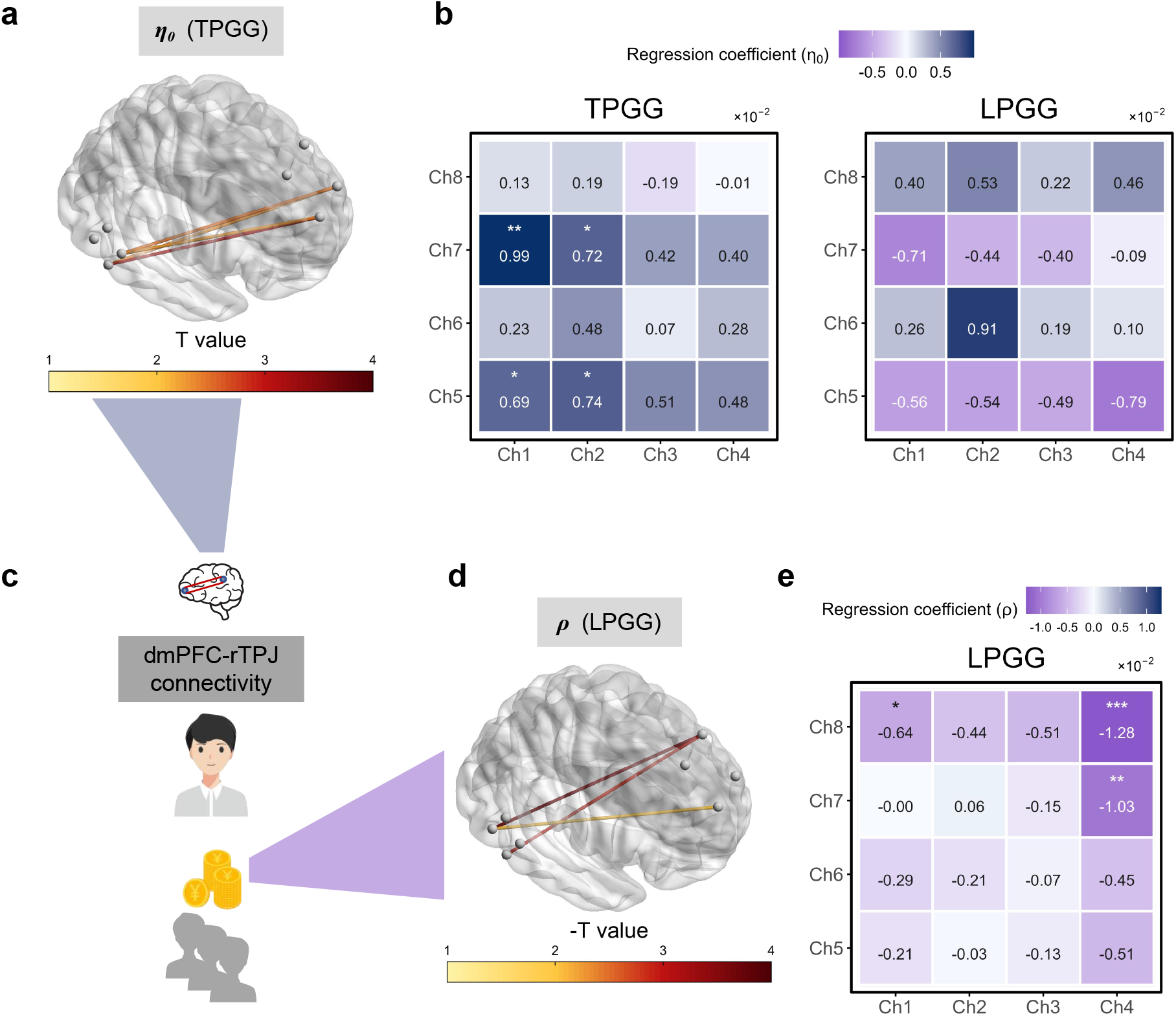
Associations between dmPFC–rTPJ functional connectivity and consequence-based social parameters. **c**, Schematic of the individual-level analyses linking dmPFC–rTPJ functional connectivity to model parameters. **a,b**, Associations between dmPFC–rTPJ connectivity and baseline consequence-based social weight η_0_. In TPGG, stronger connectivity across a subset of dmPFC–rTPJ channel pairs was positively associated with η_0_ (**a**); the corresponding regression coefficients for all 16 channel pairs are shown in **b. d,e**, Associations between dmPFC–rTPJ connectivity and context-dependent modulation ρ in LPGG. Greater *ρ* was associated with reduced connectivity across a subset of dmPFC–rTPJ channel pairs (**d**); regression coefficients for all channel pairs are shown in **e**. Lines in the brain renderings indicate channel pairs showing the corresponding associations, with colour encoding the magnitude of the *t* statistic.

### Significant inter-brain neural synchrony emerged in TPGG but not LPGG

Finally, we asked whether the distinct norms dominating the two game structures were accompanied by different strength of inter-brain neural synchrony (INS). We quantified pairwise inter-brain coherence among members of each four-person group using wavelet transform coherence (WTC), and baseline-corrected task-related coherence by subtracting resting-state coherence; the resulting pair-wise estimates were then averaged to derive group-level INS (INS; Figure 7, a). INS thus provided a complementary measure of interpersonal neural alignment during collective interaction [23].

**Fig. 7:**
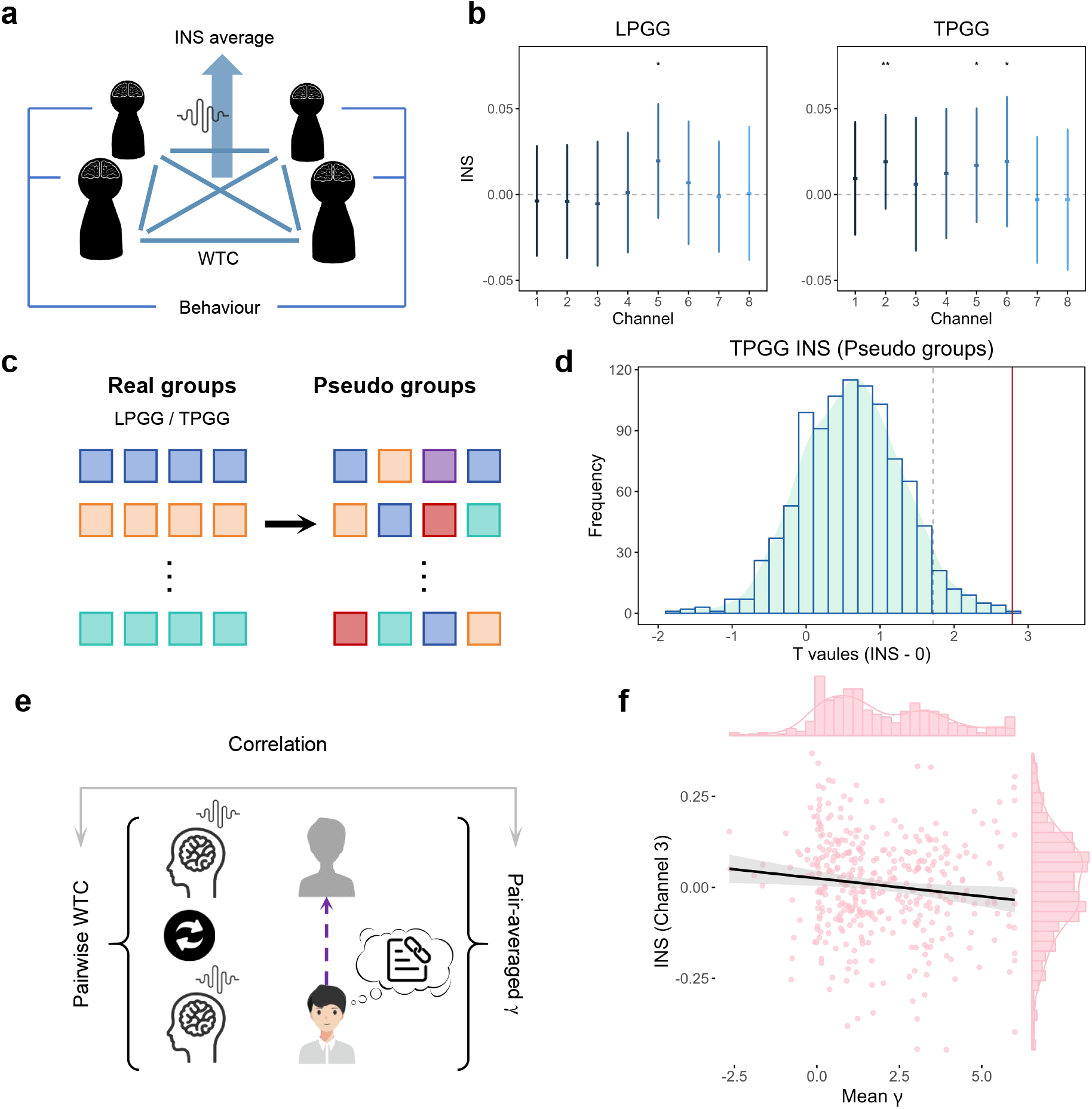
Inter-brain neural synchrony and its association with group-level conformity. **a**, Schematic of group-level inter-brain neural synchrony (INS). Wavelet transform coherence (WTC) was calculated for each of the six participant pairs within a four-person group and averaged to obtain group-level INS for each channel. **b**, Task-related INS in LPGG and TPGG. **c**, Construction of pseudo-groups for permutation analysis. Participants were reassigned within each game structure while ensuring that members of the same real group were never assigned to the same pseudo-group. **d**, Permutation test of dmPFC channel 2 synchrony in TPGG. The distribution shows test statistics obtained from 1,000 pseudo-group permutations, and the vertical red line indicates the statistic observed for the real interacting groups. **e**, Schematic of the pair-level analysis linking paired INS to pair-averaged conformity-based weight *gamma*. **f**, In TPGG, paired INS in dmPFC channels 3 was negatively associated with pair-averaged γ. Lines denote linear fits with 95% confidence intervals; marginal distributions show the corresponding variable distributions.

In LPGG, enhanced synchrony was observed only in rTPJ channel 5, whereas in TPGG it emerged in dmPFC channel 2 and rTPJ channels 5 and 6 (Figure 7, b; LPGG: channel 5, *t*_63_ = 2.78, *P* = 0.0072, Cohen’s *d* = 0.35, 95% CI [0.10, 0.60]; TPGG: channel 2, *t*_63_ = 3.05, *P* = 0.0034, Cohen’s *d* = 0.46, 95% CI [0.13, 0.63]; channel 5, *t*_63_ = 3.00, *P* = 0.0039, Cohen’s *d* = 0.38, 95% CI [0.13, 0.62]; channel 6, *t*_63_ = 2.36, *P* = 0.0216, Cohen’s *d* = 0.31, 95% CI [0.05, 0.54]; all effects survived FDR correction). To determine whether this effect reflected interaction-specific coupling rather than shared task-evoked fluctuations, we generated 1,000 pseudo-groups for each game structure by permuting participants across real groups (Figure 7, c). dmPFC channel 2 in TPGG exceeded the pseudo-group distribution (Figure 7, d; *P*_perm_ = 0.002). By contrast, rTPJ synchrony did not survive pseudo-group testing (Supplementary Figure 19), suggesting that these effects were more consistent with shared task-related fluctuations than with interaction-specific neural alignment. Together, these findings identify dmPFC synchrony as a group-level neural correlate of TPGG. This interaction-specific alignment is consistent with the stronger conformity-based norms observed in TPGG, where behavior was more persistently organized around a shared contribution standard [43–45].

Next, we examined whether INS was associated with parameter related to conformity-based norms in TPGG (Figure 7, e). The pair-averaged conformity-based weight γ was negatively associated with INS in channel 3 (Figure 7, f; β = *−*0.00989, SE = 0.00400, *t*_376_ = *−*2.47, 95% CI [*−*0.01775, *−*0.00203], *P* = 0.0138). This inverse association may reflect differences in how groups coordinate their contributions. When conformity to a shared contribution standard is weaker, group members may rely more heavily on reciprocal monitoring and complementary behavioral adjustments, rather than simply matching one another’s actions [46]. Accordingly, stronger dmPFC INS in groups with lower mean γ may reflect greater engagement of interpersonal coordination processes.

## Discussion

Our study shows that game structure reorganizes norms through which cooperation is sustained [1, 2, 5]. In LPGG, contribution behavior was more dynamically updated and more strongly shaped by others’ contributions, consistent with higher consequence-based social weight and utility [9, 47]. By contrast, TPGG showed greater stickiness to a contribution standard, consistent with stronger conformity-based weight [48, 49]. These distinct norms were accompanied by corresponding differences in regional neural activation, dmPFC–rTPJ connectivity and inter-brain neural synchrony [20, 24, 44], indicating that game structure reorganizes both the computational and neural characteristics of collective decision-making (Supplementary Table 1 and Figure 1).

The model-free behavioral results first revealed distinct contribution dynamics between two game structures [9]. Peer punishment increased contributions in both games [31, 32, 50], but its influence on contribution dynamics was more pronounced in LPGG. Contributions in LPGG increased progressively during P stage, whereas TPGG showed comparatively stable trajectories despite more severe punishment of low contributors. Conditional contribution analyses further showed that current decisions depended more strongly on individuals’ own previous contributions in TPGG, whereas LPGG showed greater imitation to peer behavior [9, 47, 51].

The computational results identified distinct norms underlying these behavioral patterns [1, 4]. LPGG was best described by the *S*-*vCS*-*C* model and TPGG was best described by the *S*-*f CS*-*C* model. Parameter estimates further showed that the baseline consequence-based social weight η_0_ was consistently higher in LPGG, whereas conformity-based weight γ was consistently greater in TPGG [2, 7, 10]. The trial-wise utility trajectories provided a dynamic view of the same dissociation. In LPGG, consequence-based social utility increased markedly when punishment became available and partially declined after punishment was removed, whereas conformity-based utility remained comparatively weak [3]. TPGG showed the opposite profile, with relatively modest consequence-based social utility but a stronger conformity-based component [49].

These results argue that the two structures shape different norms. In LPGG, contribution decisions were supported primarily by consequence-based social utility whose influence remained sensitive to social context [2, 7, 52]. In TPGG, contribution decisions were more strongly constrained by deviations from an inferred group standard [1, 10].

The neural results mirrored the computational distinction between consequence-based social norms in LPGG and more stable conformity in TPGG [20]. TPGG elicited stronger rTPJ activation, which was positively associated with contributions [19, 44, 45]. Also, dmPFC–rTPJ connectivity was positively associated with the baseline consequence-based social weight η_0_ in TPGG, whereas in LPGG connectivity was negatively associated with the context-dependent modulation ρ [19, 44]. This distinction extended to the group level [24]. Interaction-specific dmPFC inter-brain neural synchrony emerged in TPGG and survived pseudo-group permutation testing, whereas no comparable dmPFC synchrony was detected in LPGG [23]. In TPGG, linear regression revealed a negative association between the group-averaged conformity weight γ and dmPFC INS in channel 3. This inverse relationship suggests that stronger conformity to a shared contribution standard may reduce the need for ongoing interpersonal coordination, whereas weaker conformity may require greater reciprocal monitoring and complementary behavioral adjustments between group members. It high-lights the possibility that INS reflects the demands of coordinating behavior rather than the strength of conformity itself [23, 24, 46, 53].

Together, these neural findings suggest that game structure reorganized social computation across multiple neural scales [44, 45]. TPGG was characterized by stronger individual-level social-cognitive engagement and interaction-specific group alignment, whereas LPGG showed neural signatures more closely related to context-sensitive social value. The inverse association between conformity sensitivity and INS further suggests that interpersonal synchrony may reflect the need for online coordination rather than the strength of conformity itself [23, 24].

More broadly, these findings indicate that game structures can alter not only the incentives for contribution but also the normative computations through which cooperation is organized [1]. Continuous-return environments like LPGGs may favour mechanisms that preserve sensitivity to changing social context, whereas threshold environments like TPGGs may favour mechanisms that clarify and stabilize shared behavioral standards [49]. This distinction has implications for interventions aimed at sustaining collective decision-making in societal challenges such as climate change, energy sustainability and social inequality [3] and may become increasingly relevant in human–AI collective systems [54, 55]. Artificial agents designed to support cooperation may need to infer whether a group is operating in a socially responsive regime, in which recent behavior should dynamically modulate recommendations, or in a coordination regime, in which maintaining a shared behavioral standard is more important. Treating cooperation as a single behavioral objective could obscure these qualitatively different routes to collective order and lead artificial agents to intervene in ways that are poorly matched to the underlying interaction structure [54, 55].

In summary, the present study suggests that collective structure determines not merely how much individuals cooperate, but how cooperation is organized by different norms[1, 2]. LPGGs favoured flexible, context-sensitive value of the consequences of one’s actions for others, whereas TPGG interactions stabilized conformity [9, 49]. These distinct norms were expressed across behavioral dynamics, computational parameters, regional neural activation, dmPFC–rTPJ connectivity and inter-brain neural synchrony [23, 44]. Game structure therefore reshapes the behavioral and neural characteristics through which norms support collective action.

## Methods

### Participants

We performed G*power to determine the sample size [56]. To ensure adequate statistical power for detecting behavioral and neural dimension differences between LPGG and TPGG, we employed a fixed-effects ANOVA model to estimate the required sample size. For a two-group comparison, we set the effect size *f* = 0.3, significance level α = 0.05, and desired statistical power 1 *−* β = 0.9. These parameters yielded a required total sample size of 120.

We recruited 536 undergraduate participants from Beihang University. Five groups (20 participants) were excluded due to failure to arrive on time, inability to understand the procedure, or accidental withdrawal during the experiment. Ultimately, 516 participants (129 groups, 264 females, aged 17 - 25) remained for behavioral analysis. All participants received cash rewards ($12 - 20) based on experimental performance. One group was excluded from neural analysis due to fNIRS recording issues, leaving 512 participants (128 groups) for neural analysis. This met the sample size requirement estimated by G*power.

All analyzed participants were right-handed, had normal or corrected-to-normal vision, possessed basic computational skills, and completed control problems. The experiment was approved by the Beihang University Biomedical Ethics Committee (protocol no. BM20250183). The preregistration for this study is available in the OSF repository (https://osf.io/9n36j/). Written informed consent was obtained from all participants before the task. The study adhered to the Declaration of Helsinki and other relevant international ethical guidelines to ensure the safety and well-being of participants.

### Experimental procedure

The experimental procedure comprised three consecutive stages: pre-experiment assessment and instructions, the main experimental task, and a post-experiment survey. Each session lasted approximately 100 min.

### Pre-experiment assessment and instructions

On the day before the main experiment, registered participants attended a pre-experiment session at a designated site and completed a questionnaire assessing eligibility and collecting background measures for subsequent analyses (Supplementary Section Pre-experiment questionnaire). We recorded participants’ age, gender and academic background, and assessed incentivized economic preferences and social value orientation (SVO). Incentivized economic preferences were measured using a validated preference module developed in previous work [57], which elicits risk preference, patience, trust, altruism, and positive and negative reciprocity through a series of incentive-compatible decision scenarios. SVO was assessed using the SVO Slider Measure [58], which quantifies preferences for allocating resources between oneself and others and provides an index of social value orientation (Supplementary Figure 2). Participants who met the eligibility criteria then read the experimental instructions and provided written informed consent. They were explicitly informed that participation was permitted only once.

To minimize behavioral biases arising from misunderstandings of the LPGG and TPGG structures, we developed three sets of comprehension questions based on previous work [59, 60]. These questions required participants to map contribution and punishment decisions onto the corresponding payoff outcomes in each game (Supplementary Section Control questions). The experiment proceeded only after all participants had answered every comprehension question correctly.

### Main experiment

The task was framed as an investment in a virtual company and introduced through on-screen instructions accompanied by standardized audio guidance. Participants were assigned in equal numbers to one of four experimental conditions in a 2 *×* 2 design crossing game type (LPGG versus TPGG) with group composition (two high-productivity participants versus one high-productivity participant). Each participant was randomly assigned an identity code (A, B, C or D) and an individual productivity level, both of which remained fixed throughout the experiment. Identity codes were anonymized and permanently dissociated from participants’ real identities. Participants were never informed of the mapping between identity codes and the identities of other group members. Participants were then fitted with the fNIRS equipment. Before the formal task, they completed a 120-s resting-state baseline recording while remaining still. The formal task consisted of 51 decision-making rounds divided into four stages: preparation (rounds 1–6), NP1 (rounds 7–21), P (rounds 22– 36), and NP2 (rounds 37–51). The preparation, NP1 and NP2 stages shared the same structure and ended after the investment phase. By contrast, the P stage additionally included a peer-punishment phase, in which participants could incur a cost of 2 points to reduce another group member’s payoff by 6 points. To minimize end-game effects, participants were not informed of the total number of rounds, and no round counter was displayed [27]. Throughout the experiment, participants were instructed to remain silent, attend only to their own computer screen, and avoid observing other participants or their displays. To minimize motion artefacts during fNIRS recording, participants were instructed to operate the mouse with their right hand while keeping their left hand resting on the table. Mouse movements and clicks were permitted only during the investment and punishment phase, whereas participants were required to remain still during inter-trial rest periods. Keyboard use was not permitted.

#### P stage of LPGG

Each round began with a 5-s fixation period signalling the imminent onset of the trial. Participants then entered the investment phase, comprising a 5-s deliberation period followed by a 7-s response period. During the response period, they selected a contribution between 0 and 30 points using a horizontal slider marked in 5-point increments. After a 2–3-s interval, during which the system computed and recorded the round outcomes, contribution results were displayed for 10 s. The outcome screen showed productivity, contribution, weighted contribution (contribution *×* productivity), retained wealth and round payoff for each group member. To facilitate rapid interpretation, outcomes were presented using combined graphical and numerical displays, with each participant’s own information shown in the top row of each on-screen module.

Participants then entered the punishment phase. After a 5-s deliberation period, they had 7 s to indicate, using the mouse, whether to punish any of the other group members. Selecting a participant’s identity code caused the corresponding box to turn red, indicating that the target had been selected for punishment. Following a further 2–3-s interval, during which the interface displayed the number of punished targets and the total wealth spent on punishment, the final outcome of the round was presented for 7 s. This screen displayed the participant’s contribution-related payoff, any loss resulting from punishment received, and the resulting final payoff for that round. Punishment was fully anonymous: participants were never informed of the identity of those who had punished them. Consecutive rounds were separated by a 10-s rest period.

#### NP stage of LPGG

The NP stages followed the same sequence but omitted the punishment phase. After contribution outcomes had been displayed for 10 s, the final outcome screen was presented immediately for 7 s, with the punishment-loss field fixed at zero.

#### Stages of TPGG

The TPGG followed the same overall procedure as the LPGG, with the addition of a round-specific threshold. During the investment phase, the threshold for the current round was continuously displayed at the bottom of the screen throughout both the 5-s deliberation period and the 7-s contribution period. During the subsequent 10-s outcome display, the interface additionally indicated whether the group’s total weighted contribution had met or exceeded the threshold.

Thresholds were defined at three levels. The high threshold corresponded to the total weighted contribution obtained when all four group members contributed 20 of their 30 points, yielding 192 points in the two-high-productivity-participant (2HPP) condition and 160 points in the one-high-productivity-participant (1HPP) condition. The middle threshold corresponded to universal contributions of 15 points, yielding 144 points in the 2HPP condition and 120 points in the 1HPP condition. The low threshold corresponded to universal contributions of 10 points, yielding 96 points in the 2HPP condition and 80 points in the 1HPP condition. Each threshold level (high, middle and low) occurred equally often within each stage: twice during the preparation stage and five times during each of the NP1, P and NP2 stages. Threshold order was fully randomized within each stage. To introduce additional unpredictability, each threshold was independently jittered: with probability 0.25 it was increased by 8 points, with probability 0.25 it was decreased by 8 points, and otherwise it remained unchanged.

### Post-experiment survey

The post-experiment survey was designed to assess behavioral and cognitive changes over the course of the study and to obtain detailed feedback on the main task (Supplementary Section Post-experiment survey). It measured participants’ expectations about others’ contributions, perceived fit with the task across the NP and P stages, and the motivations underlying their contribution and punishment decisions.

### Behavioral data acquisition and model-free analysis

To precisely capture real-time interactions and time-stamp every experimental phase, we deployed a web-based platform that enabled four users to participate simultaneously. The application was built with a Vue.js front-end and a MySQL back-end for data capture and storage, recording each participant’s contribution and punishment choices every round. WebSocket was employed to synchronize front-end and back-end communications, enforce a consistent operation sequence across users, and log millisecond-level timestamps for every webpage transition, which were then aligned with the fNIRS neural recordings to achieve precise event marking.

### Dynamics based on round progression and aggregate punishment

To characterize model-free behavioral dynamics, we first quantified round-by-round contributions separately for LPGG and TPGG. Because no round-wise differences between high- and low-productivity participants survived FDR correction, the two productivity groups were pooled for subsequent analyses. We then tested whether punishment availability altered contribution trajectories using linear mixed-effects models, with individual contribution as the dependent variable; punishment availability (*Pun*; NP versus P), centred round number (*round*_*c*_) and their interaction as fixed effects; and participant identity as a random intercept. Significant *Pun × round*_*c*_ interactions were followed by simple-slope analyses to estimate round-wise changes in contribution separately for NP and P stages. To complement these trajectory-based analyses, we quantified aggregate punishment during the punishment stage by summing the punishment received by each participant and used linear regression to test its association with that participant’s total contribution.

### Analyses of conditional behavior

To characterize conditional behavioral dynamics, we analyzed contribution and punishment decisions using Bayesian mixed-effects models. Conditional behavior was defined as the extent to which decisions in the current round depended on behavior observed in the preceding round (Figure 2, a). For contribution decisions, we modelled each individual’s contribution in round *t*, 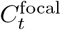, as a function of the focal individual’s own contribution in the preceding round, 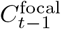, and the total contribution of the other group members, 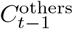 [14, 39]. The corresponding coefficients were interpreted as ***stickiness*** and ***imitation***, respectively [51]. Higher stickiness indicates greater persistence in one’s own previous contribution, whereas higher imitation indicates stronger influence by others’ previous contributions [61, 62]. We additionally included their interaction and individual-specific random intercepts and random slopes:

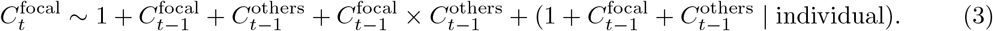

In supplementary analyses, we repeated the model using weighted contributions (contribution *×* productivity) and punishment decisions. For the TPGG, we further tested whether threshold magnitude and threshold attainment modulated these conditional contribution patterns.

### Agent-based modelling

We fitted a set of candidate models to identify the latent motives underlying individual contribution decisions. Across models, each available contribution was evaluated using different combinations of three separable utility components: self-regarding material utility, consequence-based social utility, and conformity-based utility. Rather than introducing an additional belief-learning process, we used the mean observed contributions of other group members in the immediately preceding round to calculate the current social context and shared contribution standard [4, 10].

### The components of utility

#### Self-regarding material utility

For each participant, we evaluated the seven available contribution decisions while treating the other three group members as the social reference set. In the experiment, contributions were selected from *C*_*i,t*_ *∈ {*0, 5, 10, 15, 20, 25, 30*}*. For model fitting, these seven options were rescaled to the integer action space *a*_*i,t*_ *∈ A* = *{*0, 1, 2, 3, 4, 5, 6*}*, with the mapping *C*_*i,t*_ = 5*a*_*i,t*_. For a candidate action *a ∈ A* by focal participant *f* in round *t*, the other three group members’ actions were taken from the immediately preceding round. The action profile was defined as

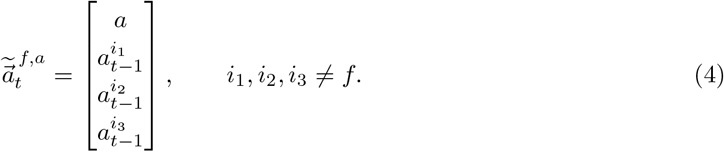

Self-regarding material utility was defined directly as the focal participant’s payoff under each candidate action:

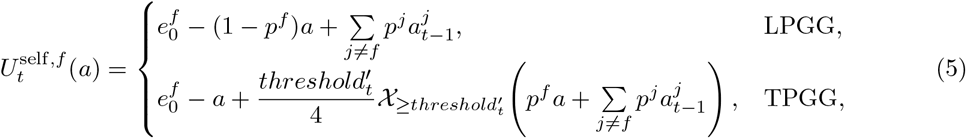

where 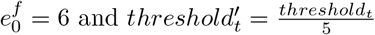.

#### Consequence-based social utility

We defined consequence-based social utility as the value assigned to how the focal participant’s action affects other group members [5]. Because higher contributions generate greater benefits for others, the social consequence of candidate action *a* was simply defined as

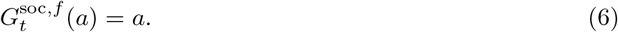

Evidence from repeated public goods games shows that cooperation is often conditional on the recent cooperative behavior of others [9, 37]. We therefore allowed the weight placed on social consequences to depend on the other group members’ contributions in the preceding round. Recent group contribution context was normalized as

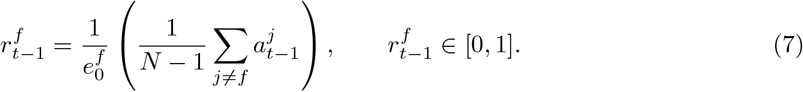

The weight assigned to social consequences was then

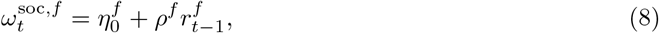

where 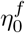 denotes the baseline weight placed on the consequences of one’s action for others, and ρ^*f*^ captures the extent to which this weight is modulated by recent group context. Consequence-based social utility was therefore

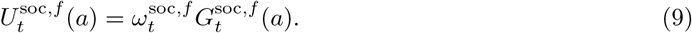

#### Conformity-based utility

We treated the recent behavior of the other group members as a social reference from which the focal participant could infer a local contribution standard [5]. The reference level was defined as the mean contribution of the other three group members in the preceding round:

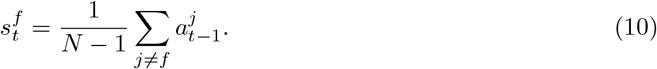

Following norm-based utility formulations [1, 4], deviations from this standard incurred a quadratic conformity cost:

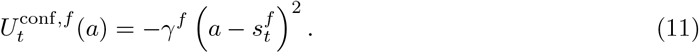

Here, γ^*f*^ captures the focal participant’s sensitivity to deviations from the inferred contribution standard.

### Utility-based models

We compared six nested candidate models constructed from three utility components: self-regarding material utility (*S*), consequence-based social utility (*CS*), and conformity-based utility (*C*). Self-regarding utility was included in all models as the common baseline. For models containing *CS*, we distinguished between a variable-weight specification (*vCS*), in which the weight assigned to social consequences was modulated by the recent contributions of others through context-dependent modulation ρ^*f*^, and a fixed-weight specification (*f CS*), in which ρ^*f*^ = 0 and the social-consequence weight remained at its baseline level 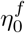.

- *S***-***vCS***-***C* **model**. This full model included all three utility components. The baseline social-consequence weight 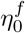 was dynamically modulated by ρ^*f*^, while γ^*f*^ captured sensitivity to deviations from the locally inferred contribution standard.
- *S***-***f CS***-***C* **model**. This model included all three utility components but assumed that consequence-based social weight remained fixed across rounds. Accordingly, ρ^*f*^ was fixed at zero, leaving 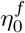 and γ^*f*^ as the social-utility parameters.
- *S***-***vCS* **model**. This model omitted conformity-based utility by fixing γ^*f*^ = 0, while retaining both the baseline social-consequence weight 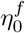 and its context-dependent modulation ρ^*f*^.
- *S***-***f CS* **model**. This model omitted conformity-based utility and assumed a stable social-consequence weight by fixing both γ^*f*^ = 0 and ρ^*f*^ = 0.
- *S***-***C* **model**. This model omitted consequence-based social utility by fixing 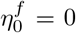 and ρ^*f*^ = 0, while retaining conformity sensitivity γ^*f*^.
- *S* **model**. This baseline model contained only self-regarding material utility, with 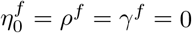.

These nested specifications allowed us to separately test whether contribution decisions were better explained by consequence-based social utility, conformity-based utility, or both. Comparisons between the *vCS* and *f CS* variants tested whether consequence-based social social weight was dynamically modulated by recent group context, whereas comparisons between models with and without *C* assessed the contribution of conformity. The *S* model provided a common material-payoff baseline.

### Model fitting and selection

Each candidate model was fitted separately for each participant and experimental stage (NP1, P and NP2), assuming a single parameter set within each stage. Parameters were estimated by maximum likelihood. For focal participant *f*, the utility of each candidate action *a ∈ A* was converted into a probability using a softmax function:

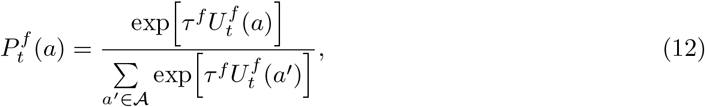

where τ ^*f*^ > 0 is the inverse-temperature parameter.

For each participant and stage, the log-likelihood of the observed contribution sequence was

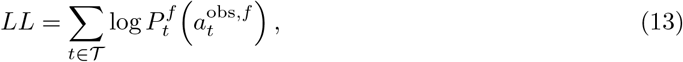

where *T* denotes the fitted rounds and 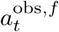 the observed action. Lagged social information, including recent group context 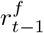 and the contrition standard 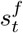, was computed directly from the other group members’ observed actions in the preceding round. For the first round of each stage, information from the immediately preceding experimental round was used; for NP1, the final preparation round provided the initial lagged observations.

Model fit was compared using the Bayesian Information Criterion (BIC),

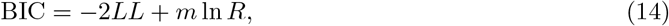

where *m* is the number of free parameters and *R* the number of fitted rounds. For each participant and stage, relative model fit was summarized as

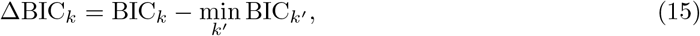

such that ΔBIC = 0 identified the best-fitting candidate model.

### Parameter recovery

We assessed whether the free parameters of the winning models could be reliably recovered from synthetic data. Parameter recovery was performed separately for the *S*-*vCS*-*C* model selected for LPGG and the *S*-*f CS*-*C* model selected for TPGG. For each participant and experimental stage, synthetic contribution sequences were generated using the individually fitted parameters and the same task structure as in the experiment. The observed contributions of the other three group members were retained to provide the one-round-lag social context, whereas the focal participant’s contribution on each round was sampled from the softmax probabilities predicted by the generating model.

We generated 100 synthetic datasets and refitted the corresponding generating model to each dataset using the same maximum-likelihood procedure applied to the empirical data. Recovery was quantified by the correspondence between the generating and re-estimated values of each free parameter, using Pearson correlations across participant-by-stage observations (Supplementary Figure 11).

### fNIRS data acquisition

Neural data from all four participants was acquired simultaneously with the ETG-4100 optical topography system (Hitachi Medical Corporation). The system samples at 10 Hz and employs near-infrared wavelengths of 695 nm and 830 nm. We used four identical customized optode probe sets, each with two sets of probes. Optode positions were localised according to the international 10-10 system using the fNIRS Optodes’ Location Decider [63]. The first probe set was positioned over the dorsal medial prefrontal cortex, with its posterior optode centered at Fz. The second probe set covered the right temporoparietal junction, aligned so that its upper-left photodiode was placed at CP6 (Supplementary Figure 15). Each probe set comprised four measurement channels—two emitters and two detectors spaced 30 mm apart. Channels 1-4 sampled the medial prefrontal cortex, whereas channels 5-8 targeted the right temporoparietal junction [64]. Before every session, all optode arrays were inspected and repositioned as needed to ensure consistent placement both within and across participant groups. Following the modified Beer-Lambert law, concentration changes in HbO and HbR were extracted from each channel. Because HbO is the most sensitive haemodynamic variable and shows a strong positive correlation with the BOLD signal, only HbO concentration changes were analyzed.

### fNIRS data processing

#### Data preprocessing

We focus on three neural scales: (1) neural activation; (2) functional connectivity; and (3) interbrain neural synchrony. The raw fNIRS data were first detrended using a DCT high-pass filter with a 128-s cut-off, and temporal autocorrelation was corrected via a pre-colouring approach whose filter shape is HRF transfer function [65].

#### Neural activation

We quantified task-related activation from the preprocessed oxyhaemoglobin (HbO) signals during contribution decisions. Each trial contained an initial 10-s baseline interval, corresponding to the waiting or outcome-feedback period, followed by the decision period during which participants considered and selected their actions (Figure 5, a). For each channel and trial, HbO signals during the decision period were standardized relative to the mean and standard deviation of the corresponding baseline interval. Task-related activation was defined as the mean baseline-standardized HbO signal during the decision period.

We first compared channel-wise activation between LPGG and TPGG to identify game-structure-dependent neural responses, and subsequently examined activation across NP1, P and NP2 in channels showing robust between-game differences. To link neural activation to individual behavior and computational processes, participant-level mean activation was further related to mean contributions and to the fitted inverse-temperature parameter τ.

#### Neural functional connectivity

We next examined functional coupling between the dmPFC and rTPJ during contribution decisions. Wavelet transform coherence (WTC) was computed between each dmPFC channel (channels 1–4) and each rTPJ channel (channels 5–8), yielding 16 within-brain channel pairs. Coherence estimates were Fisher *z*-transformed before statistical analysis.

For each participant, we obtained the mean connectivity strength of each dmPFC–rTPJ pair and related these values to the fitted normative parameters. Analyses focused on the baseline consequence-based social weight η_0_ and the context-dependent modulation ρ. Linear regressions were performed separately for LPGG and TPGG for each channel pair.

#### Inter-brain neural synchrony

We quantified inter-brain neural synchrony (INS) from the preprocessed HbO signals using wavelet transform coherence (WTC). For each participant and channel pair, task-related INS was defined as the increase in Fisher *z*-transformed coherence during the contribution decision relative to the pre-experiment resting-state baseline.

We first identified the frequency of interest (FOI) from the task–rest coherence difference averaged across participant pairs, channels and experimental conditions over the 142 sampled frequency bins. The strongest task-related increase occurred between 0.1219 and 0.1536 Hz, corresponding to oscillatory periods of approximately 6.5–8.2 s (Supplementary Figure 17). Subsequent INS analyses were therefore restricted to this frequency range.

For each four-person group, INS within the FOI was first averaged for each of the six possible participant pairs and then averaged across pairs to obtain a group-level INS estimate for each channel (Figure 7, a). We tested whether task-related INS differed from zero within each game structure.

To determine whether the observed synchrony reflected interaction-specific neural coupling rather than neural fluctuations shared across individuals performing the same task, we conducted pseudo-group permutation analyses. Participants were reassigned to pseudo-groups within the same game structure, with the constraint that no pseudo-group contained members of the same real interacting group. For each of 1,000 permutations, INS was recomputed using the same procedure applied to the real groups, generating empirical null distributions for within-game synchrony (Figure 7, c). Permutation significance was calculated as

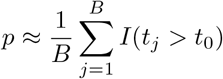

where *I* is the indicator function.

Finally, we examined whether inter-brain synchrony varied with model-derived conformity sensitivity. For each participant pair, the fitted conformity-based weight γ was averaged across the two individuals to obtain a pair-level conformity index. For each channel, INS was calculated as described above and regressed on pair-averaged γ using ordinary regression in TPGG (Figure 7, e).

#### Analysis plan

All statistical analyses and visualizations were performed in R (v4.3.1) and MATLAB (R2022a).

For behavioral analyses, Bayesian multilevel regression models were fitted using *brms* (v2.22.0) [66]. Conditional contribution patterns were analyzed using Gaussian multilevel models, whereas conditional punishment was modeled using a Bernoulli likelihood. Continuous predictors and dependent variables were standardized to zero mean and unit variance before fitting; binary punishment variables were left unstandardized. Models included participant-specific random intercepts and, where appropriate, random slopes for the corresponding predictors to account for repeated observations and individual heterogeneity. Posterior estimates are reported as posterior means with 95% highest-density intervals (HDIs), calculated using *bayestestR* (v0.15.2). Models were estimated using 10 Markov chain Monte Carlo chains, each with 1,000 warm-up and 3,000 post-warm-up iterations. Convergence was assessed using 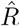 together with bulk and tail effective sample sizes; all 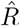 values were below 1.05.

Computational models were fitted separately for each participant and experimental stage by maximum likelihood in MATLAB. Free parameters were estimated by minimizing the negative log-likelihood using fmincon. To reduce sensitivity to local minima, optimization was repeated from 500 independently initialized starting points, and the solution with the lowest negative log-likelihood was retained. Candidate models were compared using the Bayesian Information Criterion (BIC).

For neural analyses, fNIRS data were preprocessed in MATLAB using routines based on NIRS-SPM [65]. Individual-level analyses quantified task-related HbO activation and dmPFC–rTPJ functional connectivity, whereas group-level analyses quantified inter-brain neural synchrony using wavelet transform coherence (WTC). Because the experiment was prolonged, the channel coupling showed a gradual decline across rounds (Supplement Figure). To minimize potential time-on-task effects, including fatigue and habituation, all WTC-based analyses were therefore restricted to the NP1 stage (Supplementary Figure 18). Accordingly, functional connectivity and INS analyses used only neural responses, behavioral measures and model-derived parameters from NP1 stage. Neural measures were then compared between LPGG and TPGG, while multiple comparisons across channels or channel pairs were controlled using the false-discovery rate (FDR). Furthermore, the processed neural results were related to the corresponding behavioral measures or model-derived parameters using linear regression.

## Supporting information

Supplementary Methods and Results

## Data availability

Behavioral data and processed multidimensional functional near-infrared spectroscopy (fNIRS) data from the four-person groups are publicly available on the Open Science Framework (OSF) at (https://osf.io/9n36j/).

## Code availability

The code used to reproduce the statistical analyses is publicly available on the Open Science Framework (OSF) at (https://osf.io/9n36j/).

## Acknowledgements

This work is supported by Program of National Natural Science Foundation of China (12425114, 12671643, 12501702, 12526528, 62441617), Taishan Scholars Program of Shandong Province (tstp20230635), Key R&D Program of Shandong Province (2026CXPT283), the Fundamental Research Funds for the Central Universities, the Opening Project of the State Key Laboratory of General Artificial Intelligence(Project NSKLAGI2025OP16), the special Funding for the ‘Case-by-Case-Introduction of Top Talent (Teams)’, and Beijing Advanced Innovation Center for Future Blockchain and Privacy Computing.

## Author contributions

Y.L. and X.W. conceived the project and designed the experiments. Y.L., X.W., W.D. and L.L. organized the experiments and collected the data. Y.L., X.W., W.D. and L.L. analyzed the data and prepared the figures. Y.L. and X.W. developed the computational model and interpreted the results. Y.L., X.W., L.L., X.C. and S.T. wrote the original and final version of the manuscript. X.W., S.T. and H.Z. funded and supervised the project.

