## Supplementary Methods and Results for "Game structure shapes individuals’ norms in collective decision-making"

### Supplementary Methods and Results 1

#### Experimental design details

We framed the task as an investment scenario in which participants allocated part of their endowment to a virtual corporation. Individual investments were transformed according to each participant's productivity, and the resulting corporate wealth corresponded to the return generated from the public pool in the public goods game. The peer-punishment mechanism was explained in detail before the experiment. Participants were informed that punishment would be available in some rounds, but the specific rounds in which punishment was available were not disclosed in advance (Supplementary Figure 2).

Before the main task, we measured incentivized economic preferences and social value orientation (SVO) (Supplementary Figure 3). Economic-preference measures showed broad variation across participants, indicating substantial heterogeneity in the sample. SVO scores were concentrated predominantly in the prosocial region of the payoff plane.

#### Nash equilibrium

Our comparison of the LPGG and TPGG was based on a four-person public goods game. In LPGG, no threshold was imposed. Instead, all contributions were pooled, multiplied by the corresponding productivity, and then redistributed equally across group members. Under the linear payoff structure, the one-shot game has a unique Nash equilibrium: each participant contributes nothing, such that $C_i = 0$ for all individuals.

In TPGG, by contrast, each round was associated with a threshold, and collective returns were generated only when the total weighted contribution reached that threshold. For the equilibrium analysis, we did not incorporate the threshold jitter. Although the threshold structure created the possibility that positive contributions could be payoff-maximizing when the group coordinated successfully, zero contribution ($C_i = 0$) remained a Nash equilibrium, as unilateral contribution was always risky when others failed to contribute sufficiently [1]. Additional pure-strategy Nash equilibria could arise when the group exactly reached the threshold, because any unilateral reduction in contribution would then cause the group to miss the threshold and lower the deviating individual's payoff. Specifically, coordinated pure-strategy equilibria may arise when the group exactly reaches the threshold,

$$\sum_{j=1}^{4} p_j C_j = threshold$$

The analysis yields the expected contrast between the two game structures. In LPGG, equilibrium predictions are unambiguously free-riding. In TPGG, by contrast, equilibrium predictions are mixed: zero contribution remains stable, but additional coordinated equilibria can emerge when the threshold can be met exactly at sufficiently low individual cost. Thus, relative to LPGG, the TPGG introduces the possibility of stable coordination around shared threshold, especially at low- and middle-level thresholds, while high-level thresholds imply greater coordination risk and make such equilibria less likely to be behaviorally realized.

From an interdependence perspective, the two game structures instantiate qualitatively different social situations rather than merely different payoff structures [2, 3]. In the LPGG, the mapping from contribution to collective return is continuous, such that each additional unit of contribution directly pits individual interest against collective welfare; in social interdependence terms, this structure is dominated by conflict, because actors can always improve their own immediate outcome by withholding resources even though the group would benefit from broader cooperation. By contrast, the TPGG introduces a shared provision point, so that the consequences of one's contribution depend more strongly on whether others coordinate around the same target; this shifts the interdependence structure towards behavioral control and mutual dependence, making success contingent on joint alignment rather than on isolated marginal gains. More broadly, recent interdependence theory argues that cooperation and punishment are shaped by inferred mutual dependence, power and conflict [4], and that higher perceived mutual dependence promotes both cooperation and direct responses to norm violation. Accordingly, the threshold structure used here can be understood as increasing perceived mutual dependence around a salient collective target, whereas the linear structure preserves a more continuously conflictual environment that favors flexible, individually responsive adjustment.

#### Threshold-dependent behavioral patterns in TPGG

We next examined whether reaching the threshold shaped behavior in TPGG [5]. Peer punishment increased the probability of threshold attainment across rounds and groups, with larger effects at the middle and high thresholds than at the low threshold. At the round level, threshold attainment was higher during the P stage than during the NP stages for the high and middle thresholds, but not for the low threshold (Supplementary Figure 4, b; Wilcoxon signed-rank tests: high threshold, $P = 0.0048$, $r = 0.744$, 95% CI [0.46, 0.85]; middle threshold, $P = 0.0233$, $r = 0.601$, 95% CI [0.21, 0.84]; low threshold, $P = 0.0755$, $r = 0.475$, 95% CI [0.04, 0.80]).

We further quantified group-level success as the proportion of rounds in which each group reached the threshold. Consistent with the round-level analysis, the increase in success rate during the P stage was larger at the middle and high thresholds and weaker at the low threshold (Supplementary Figure 4, c; high threshold, $r = 0.123$, 95% CI [0.01, 0.28]; middle threshold, $r = 0.125$, 95% CI [0.01, 0.27]; low threshold, $r = 0.118$, 95% CI [0.01, 0.26]).

#### Anti-social punishment

We next classified dyadic punishment events according to whether they targeted free riding or constituted antisocial punishment. For each punishment event, we defined the contribution deviation as the punisher's contribution minus the recipient's contribution. Positive deviations were classified as punishment of free riding, whereas non-positive deviations were classified as antisocial punishment [6, 7].

The frequency distribution of punishment events across contribution deviations is shown in Supplementary Figure 5. The proportion of antisocial punishment was lower in LPGG than in TPGG (LPGG: $\theta = 0.4836$; TPGG: $\theta = 0.5913$). Within TPGG, antisocial punishment was more prevalent when the group failed to reach the threshold than when the threshold was reached (not reaching threshold: $\theta = 0.6249$; reaching threshold: $\theta = 0.5913$).

#### Conditional contribution and punishment patterns and contribution dynamics across rounds

For the conditional punishment analysis, we modelled whether participant $i$ punished participant $j$ in round $t$, $Pun_t^{i\to j}$, as a binary outcome, with $Pun_t^{i\to j} = 1$ indicating punishment and $Pun_t^{i\to j} = 0$ otherwise. Punishment was modelled as a function of the recipient's current contribution, $C_t^j$, the punisher's own current contribution, $C_t^i$, and the punishment received by the punisher in the preceding round, $Pun_{t-1}^{G\to i}$ (Supplementary Figure 6, a). The corresponding coefficients captured *recipient-contribution sensitivity*, *self-contribution sensitivity*, and *retribution*, respectively. We additionally included all two-way interactions and the three-way interaction among these predictors. The model was fitted using Bayesian logistic mixed-effects regression, with participant-specific random intercepts and random slopes for the three main effects.

$$\begin{aligned} Pun_t^{i\to j} \sim\, & C_t^j * C_t^i * Pun_{t-1}^{G\to i} \\ & + \left(1 + C_t^j + C_t^i + Pun_{t-1}^{G\to i} \mid \text{individual}\right). \end{aligned} \tag{1}$$

where

$$Pun_t^{i\to j} = \begin{cases} 1, & \text{if participant } i \text{ punished participant } j \text{ in round } t, \\ 0, & \text{otherwise.} \end{cases} \tag{2}$$

First, retribution was stronger in TPGG than in LPGG (Supplementary Figure 6, b; LPGG: $\beta = 0.45$, 95% CI [0.32, 0.58]; TPGG: $\beta = 0.76$, 95% CI [0.60, 0.93]). This pattern is consistent with the higher prevalence of antisocial punishment in TPGG, suggesting that punishment decisions were more strongly shaped by retaliation after having been punished. Second, self-contribution sensitivity was also higher in TPGG (LPGG: $\beta = 0.43$, 95% CI [0.17, 0.70]; TPGG: $\beta = 0.69$, 95% CI [0.40, 0.90]), whereas recipient-contribution sensitivity was stronger in LPGG, as reflected by the larger negative coefficient (LPGG: $\beta = -2.29$, 95% CI [$-2.56$, $-2.04$]; TPGG: $\beta = -1.90$, 95% CI [$-2.19$, $-1.64$]). Together, these patterns suggest that punishment in TPGG was more strongly associated with the punisher's own contribution and prior punishment experience, whereas punishment in LPGG was more selectively directed towards low-contributing recipients.

Across game structures and stages, participant-specific random slopes for stickiness and imitation were negatively correlated, indicating a consistent trade-off between persistence in one's own previous behavior and responsiveness to others' contributions (Supplementary Figure 7, a, b; LPGG: $r^{\text{NP}}_{\text{random}} = -0.73$, 95% CI [$-0.85$, $-0.59$]; $r^{\text{P}}_{\text{random}} = -0.78$, 95% CI [$-0.92$, $-0.61$]; TPGG: $r^{\text{NP}}_{\text{random}} = -0.87$, 95% CI [$-0.93$, $-0.80$]; $r^{\text{P}}_{\text{random}} = -0.79$, 95% CI [$-0.88$, $-0.67$]). Random slopes for recipient-contribution and self-contribution sensitivity were also negatively correlated in both LPGG and TPGG (Supplementary Figure 7, c; LPGG: $r_{\text{random}} = -0.63$, 95% CI [$-0.77$, $-0.45$]; TPGG: $r_{\text{random}} = -0.59$, 95% CI [$-0.74$, $-0.40$]), suggesting a corresponding trade-off between punishment guided by the recipient's behavior and punishment associated with the punisher's own contribution. Repeating the conditional contribution analysis using productivity-weighted contributions (contribution × productivity) yielded qualitatively similar patterns in both LPGG and TPGG (Supplementary Figure 8).

Beyond conditional contribution patterns, we also examined round-to-round contribution dynamics, defined as $\Delta C_t^{\text{focal}} = C_{t+1}^{\text{focal}} - C_t^{\text{focal}}$ (Supplementary Figure 9, a). Changes in the focal participant's contribution were modelled as a function of changes in others' contributions and, during the P stage, punishment received in the preceding round. Punishment was positively associated with subsequent contribution change, indicating that participants increased their contributions after being punished. The $\Delta_{\text{imitation}}$ coefficient was consistently larger in LPGG than in TPGG across both NP and P stages (Supplementary Figure 9, b, c), and the punishment effect was likewise stronger in LPGG during the P stage. These results extend the conditional contribution analysis by showing that LPGG elicited greater dynamic adjustment to both peer behavior and punishment, whereas TPGG was characterized by more persistent contribution changes.

#### Context-dependent modulation

To further characterize the context dependence of consequence-based social value in LPGG, we examined the modulation $\rho$ across stages (Supplementary Figure 10, a). Mean $\rho$ increased across the experiment, from negative values in NP1 ($M = -0.538$, SD = 2.780) to approximately zero during P ($M = -0.003$, SD = 2.706) and positive values in NP2 ($M = 0.376$, SD = 3.358), with substantial individual heterogeneity at each stage. This pattern indicates a progressive shift in how recent group cooperation modulated consequence-based social value.

Decomposing consequence-based social utility into its baseline and context-dependent components further illustrated this dynamic modulation (Supplementary Figure 10, b). The context-dependent component increased when punishment became available and remained dynamically sensitive to recent group behavior, contributing to the marked increase in total consequence-based social utility during P. Following punishment removal, total social utility and baseline social value declined, while the relative contribution of context-dependent value remained pronounced.

#### Agent-based model validation

We evaluated the computational framework using parameter recovery, model recovery and forward simulation. For parameter recovery, 100 synthetic decision sequences–each comprising all stages for each participant were generated from the winning models using participant-specific fitted parameters. For each participant and experimental stage, the observed one-round-lag social context from the corresponding preceding experimental stage was used for initialization, while focal decisions were sampled from the model-predicted softmax distribution. The winning model was then re-fitted to each synthetic dataset, and recovery was quantified by Pearson correlations between the generating and recovered parameter estimates (Supplementary Figure 11). This procedure assessed whether the parameters governing self-regarding, consequence-based social and conformity-based utilities could be reliably identified from the available contribution data.

We next examined model identifiability across all six candidate models ($S$-$vCS$-$C$, $S$-$fCS$-$C$, $S$-$vCS$, $S$-$fCS$, $S$-$C$ and $S$). Synthetic datasets were generated separately from each candidate model using the corresponding participant- and stage-specific fitted parameters and the same lagged social context as in the empirical analysis. Each synthetic dataset was subsequently fitted with all six models, and the recovered model was defined as the model with the lowest BIC. Model recovery was summarized using confusion matrices. Recovery was perfect in LPGG (Supplementary Figure 12, a). In TPGG, misclassification was restricted primarily to the distinction between variable and fixed consequence-based social weight, whereas the presence or absence of the conformity component was preserved (Supplementary Figure 12, b). These results indicate that the broader model families were distinguishable, although TPGG provided more limited information for separating variable- from fixed-weight consequence-based social value.

Finally, we assessed generative adequacy using forward simulations based on the winning model for each game structure: $S$-$vCS$-$C$ for LPGG and $S$-$fCS$-$C$ for TPGG. Simulated groups reproduced the experimental productivity composition, and each agent selected among the seven contribution options according to the corresponding softmax probabilities. Recent social context and the conformity standard were updated recursively from the simulated contributions of the other group members in the preceding round, without introducing an additional belief-learning process. Simulations followed the experimental stage structure, and repeated runs were averaged to obtain model-generated contribution trajectories. These trajectories were compared with the observed behavior of high- and low-productivity participants to assess whether the inferred norms were sufficient to reproduce the characteristic contribution dynamics of LPGG and TPGG.

#### Model evolution and steady state

We next examined how the parameters of the winning models jointly shaped steady-state contribution behavior in LPGG and TPGG. To isolate the intrinsic effects of the parameters from the observed behavioral trajectories, we performed out-of-sample agent-based simulations without using empirical contribution sequences. Each simulation reproduced the four-person group structure of the experiment, with all agents assigned identical parameter values so that the contribution dynamics reflected the parameter combination under investigation rather than between-agent heterogeneity.

For LPGG, we systematically varied the baseline consequence-based social weight $\eta_0$, the context-dependent modulation $\rho$ and conformity-based weight $\gamma$. For TPGG, we varied $\eta_0$ and $\gamma$ separately across the low-, middle- and high-threshold conditions. Inverse-temperature parameter *tau* was fixed at their mean fitted values in the corresponding structure. Each simulation was initialized with two rounds of contributions randomly sampled from the experimental action space, which provided the initial social context and conformity reference.

For each parameter combination, four agents repeatedly selected among the seven contribution options according to the corresponding winning model and its softmax probabilities. The social context and conformity standard were updated recursively from the simulated contributions in the preceding round. Simulations were run for 100 rounds, which was sufficient for contribution behavior to approach a stable regime, and each parameter combination was repeated 100 times. Steady-state contribution rates were obtained by averaging across simulation runs. The analyses characterize how consequence-based social and conformity-based motives determine the long-run contribution regime in each game structure.

#### The *S-vCS-C* model of LPGG

The steady-state contribution regimes generated by the winning *S-vCS-C* model in LPGG are shown in Supplementary Figure 13. We systematically varied the baseline consequence-based social weight $\eta_0$, the context-dependent modulation $\rho$ and conformity-based weight $\gamma$, while fixing the inverse-temperature parameter at its mean fitted value ($\tau = 7.827$). The simulations revealed sharply differentiated contribution regimes across the parameter space, indicating that long-run contribution in LPGG depended on the joint configuration of consequence-based social and conformity-based value rather than on any single parameter in isolation.

Across values of $\rho$, higher $\eta_0$ generally enlarged the region associated with high steady-state contribution, consistent with a stronger baseline value of actions that benefit the group. The effect of $\rho$ depended on the surrounding parameter regime: because $\rho$ determines how strongly recent group cooperation modulates consequence-based social value, changing $\rho$ shifted the boundaries separating low- and high-contribution states. Conformity sensitivity $\gamma$ further reshaped these boundaries by determining the extent to which agents remained close to the contribution reference established by the group. Consequently, the same level of baseline social value could support markedly different steady states depending on how strongly social value was modulated by recent group behavior and how strongly agents responded to deviations from the local contribution reference.

These steady-state patterns were qualitatively similar across productivity types and group compositions (Supplementary Figure 13, a–d). Although high- and low-productivity agents, and groups containing one versus two high-productivity participants, differed modestly in the precise boundaries between contribution regimes, the overall structure of the parameter landscape remained stable. In all cases, steady-state contributions were governed primarily by the joint effects of $\eta_0$, $\rho$ and $\gamma$, indicating that the qualitative influence of consequence-based social and conformity-based value was robust to differences in productivity and group composition.

#### The $S$-$fCS$-$C$ model of TPGG

The steady-state contribution regimes generated by the winning $S$-$fCS$-$C$ model in TPGG are shown in Supplementary Figure 14. We systematically varied the baseline consequence-based social weight $\eta_0$ and conformity sensitivity $\gamma$, while fixing the inverse-temperature parameter at its mean fitted value ($\tau = 2.689$). Simulations were conducted separately for low-, middle- and high-threshold conditions and for groups containing one or two high-productivity participants. Across these conditions, the parameter space was partitioned into distinct contribution regimes, indicating that long-run behavior in TPGG depended jointly on baseline social value, conformity sensitivity and the prevailing threshold.

Across threshold levels, higher $\eta_0$ generally expanded the region associated with high contribution, whereas the effect of $\gamma$ depended on the contribution reference established by the group. The resulting parameter landscapes showed relatively sharp boundaries between low- and high-contribution states, particularly at the middle and high thresholds. This pattern is consistent with the coordination structure of TPGG: when the threshold is more demanding, small changes in social value or conformity can move the group between qualitatively different contribution regimes rather than producing only gradual changes in contribution.

The qualitative structure of these steady-state landscapes was similar across productivity classes and group compositions (Supplementary Figure 13a–d). Differences between high- and low-productivity agents, and between groups containing one versus two high-productivity participants, were reflected mainly in the precise locations of regime boundaries rather than in the overall organization of the parameter space. Likewise, increasing the threshold shifted these boundaries but preserved the general interaction between $\eta_0$ and $\gamma$. Together, these simulations indicate that steady-state behavior in TPGG is characterized by comparatively discrete coordination regimes whose stability is jointly determined by baseline consequence-based social value and conformity.

### Supplementary Methods and Results 2

#### Additional results of neural activation

Although neural activation during the contribution phase was consistently higher in TPGG than in LPGG, this between-game difference was no longer significant during the punishment phase after FDR correction (Supplementary Figure 16; Ch 5: $F_{1,506} = 0.82$, $P = 0.365$, $\eta_p^2 = 0.0016$, 95% CI [0.0000, 0.0173]; Ch 6: $F_{1,496} = 2.18$, $P = 0.141$, $\eta_p^2 = 0.0044$, 95% CI [0.0000, 0.0243]; Ch 7: $F_{1,505} = 0.00$, $P = 0.955$, $\eta_p^2 = 0.0000$, 95% CI [0.0000, 0.0000]; Ch 8: $F_{1,503} = 4.33$, $P = 0.038$, $\eta_p^2 = 0.0085$, 95% CI [0.0002, 0.0305]). This attenuation suggests that punishment recruited additional social-cognitive processing in LPGG, thereby reducing the activation difference between game structures.

Consistent with this interpretation, activation in LPGG was significantly higher during the punishment phase than during the contribution phase in channels 5, 6 and 8 (paired $t$-tests: Ch 5: $t_{255} = 3.75$, $P = 0.00022$, Cohen's $d = 0.39$, 95% CI [0.18, 0.60]; Ch 6: $t_{255} = 4.91$, $P < 0.001$, Cohen's $d = 0.51$, 95% CI [0.29, 0.73]; Ch 8: $t_{255} = 8.83$, $P < 0.001$, Cohen's $d = 0.88$, 95% CI [0.65, 1.11]; all effects survived FDR correction). These results suggest that punishment particularly enhanced rTPJ engagement in LPGG, consistent with increased processing of socially relevant information during sanctioning decisions [8].

#### Additional results of INS

We observed a progressive decline in INS across rounds in both LPGG and TPGG (Supplementary Figure 18). Focusing on dmPFC channel 2 and rTPJ channel 5, INS decreased significantly over time in both game structures (Ch 2, LPGG: $F_{1,49} = 5.01$, $P = 0.0297$, $\eta_p^2 = 0.093$; Ch 2, TPGG: $F_{1,49} = 16.55$, $P = 0.00017$, $\eta_p^2 = 0.253$; Ch 5, LPGG: $F_{1,49} = 7.19$, $P = 0.00997$, $\eta_p^2 = 0.128$; Ch 5, TPGG: $F_{1,49} = 36.21$, $P < 0.001$, $\eta_p^2 = 0.425$). Similar declining trends were observed across the remaining channels (Supplementary Figure 18, c). This temporal decrease may reflect progressive stabilization of group interaction and individual behavioral routines, thereby reducing the need for continuous inter-brain alignment. Time-on-task effects, including declining attention or fatigue over the course of the experiment, may also have contributed to this pattern.

Although task-related INS in several rTPJ channels was significantly greater than zero, these effects did not survive pseudo-group permutation testing (Supplementary Figure 19; LPGG Ch 5: $P_{\text{perm}} = 0.482$; TPGG Ch 5: $P_{\text{perm}} = 0.335$; TPGG Ch 6: $P_{\text{perm}} = 0.599$). Thus, the observed rTPJ synchrony was more consistent with neural fluctuations shared across participants performing the same task than with interaction-specific interpersonal coupling. Given the established involvement of rTPJ in representing socially relevant information and others' behavior, similar task demands across group members may have produced comparable temporal responses without requiring interaction-specific neural alignment [9, 10].

**Table 1**: Multi-scale summary of different norms between LPGG and TPGG

| **Domain** | **LPGG** | **TPGG** | **Interpretation** |
|---|---|---|---|
| **Game structure** | Linear returns; persistent conflict between private and collective interests | Returns depend on reaching a predefined threshold | TPGG adds a target-based coordination demand to the pure dilemma |
| **Behavioral dynamics** | Higher flexibility, more influenced by others' behavior | Higher stickiness, stronger consensus and stronger punishment enforcement | LPGG promotes flexible behavioral adjustment, whereas TPGG promotes consensus |
| **Agent-based models** | Best captured by a varying-weight $S$-$vCS$-$C$ model; higher baseline consequence-based social weight $\eta_0$ | Best captured by a fixed-weight $S$-$fCS$-$C$ model; large conformity-based weight $\rho$ | Consequence-based social utility varies with recent context in LPGG while stronger conformity-based alignment in TPGG |
| **Individual-level neural characteristics** | Functional connectivity between dmPFC and rTPJ was negatively associated with context-dependent modulation $\rho$ | Higher neural activation, functional connectivity between dmPFC and rTPJ was positively associated with baseline consequence-based social weight $\eta_0$ | dmPFC was recruited more strongly in TPGG, while game structure reorganized how social-cognitive network related to normative computation |
| **Group-level neural characteristics** | – | Significant INS in dmPFC | Neural alignment is consistent with the stronger conformity-based norms observed in TPGG |
| **Overall profile** | Consequence-based social norms dominate collective decision-making | Conformity-based norms dominate collective decision-making | |

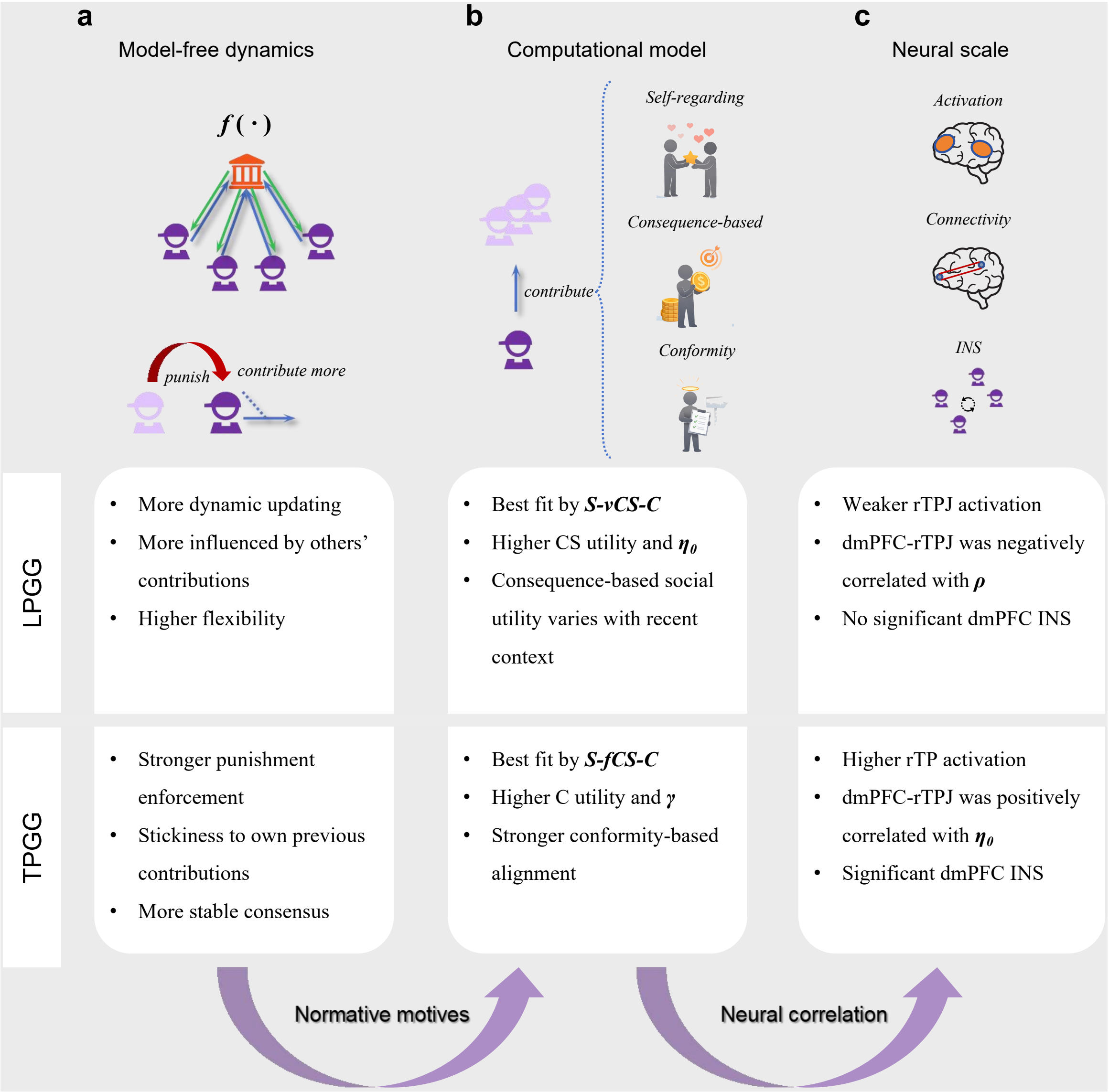

a
Model-free dynamics
f(·)
punish
contribute more
b
Computational model
Self-regarding
Consequence-based
Conformity
contribute
c
Neural scale
Activation
Connectivity
INS
LPGG
More dynamic updating
More influenced by others' contributions
Higher flexibility
Best fit by S-vCS-C
Higher CS utility and $\eta_0$
Consequence-based social utility varies with recent context
Weaker rTPJ activation
dmPFC-rTPJ was negatively correlated with $\rho$
No significant dmPFC INS
TPGG
Stronger punishment enforcement
Stickiness to own previous contributions
More stable consensus
Best fit by S-fCS-C
Higher C utility and $\gamma$
Stronger conformity-based alignment
Higher rTP activation
dmPFC-rTPJ was positively correlated with $\eta_0$
Significant dmPFC INS
Normative motives
Neural correlation

**Supplementary Fig. 1**: **Multi-scale summary of structure-dependent norms in collective decision-making.** **a**, Model-free dynamics. LPGG showed more dynamic contribution updating, greater influenced by others and higher behavioral flexibility, whereas TPGG showed stronger punishment enforcement, greater stickiness to previous decisions and more stable consensus. **b**, Agent-based modelling. LPGG was best described by the $S$-$vCS$-$C$ model, with greater consequence-based social utility and a social weight that varied with recent context. TPGG was best described by the $S$-$fCS$-$C$ model, with stronger conformity-based utility and greater conformity-based weight. **c**, Neural characteristics. LPGG showed weaker rTPJ activation, a negative association between dmPFC–rTPJ connectivity and the context-modulation $\rho$, and no significant dmPFC inter-brain synchrony (INS). TPGG showed stronger rTPJ activation, a positive association between dmPFC–rTPJ connectivity and the baseline consequence-based social weight $\eta_0$, and significant dmPFC INS. Together, these findings indicate that game structure reorganizes norms across behavioral, computational and neural scales.

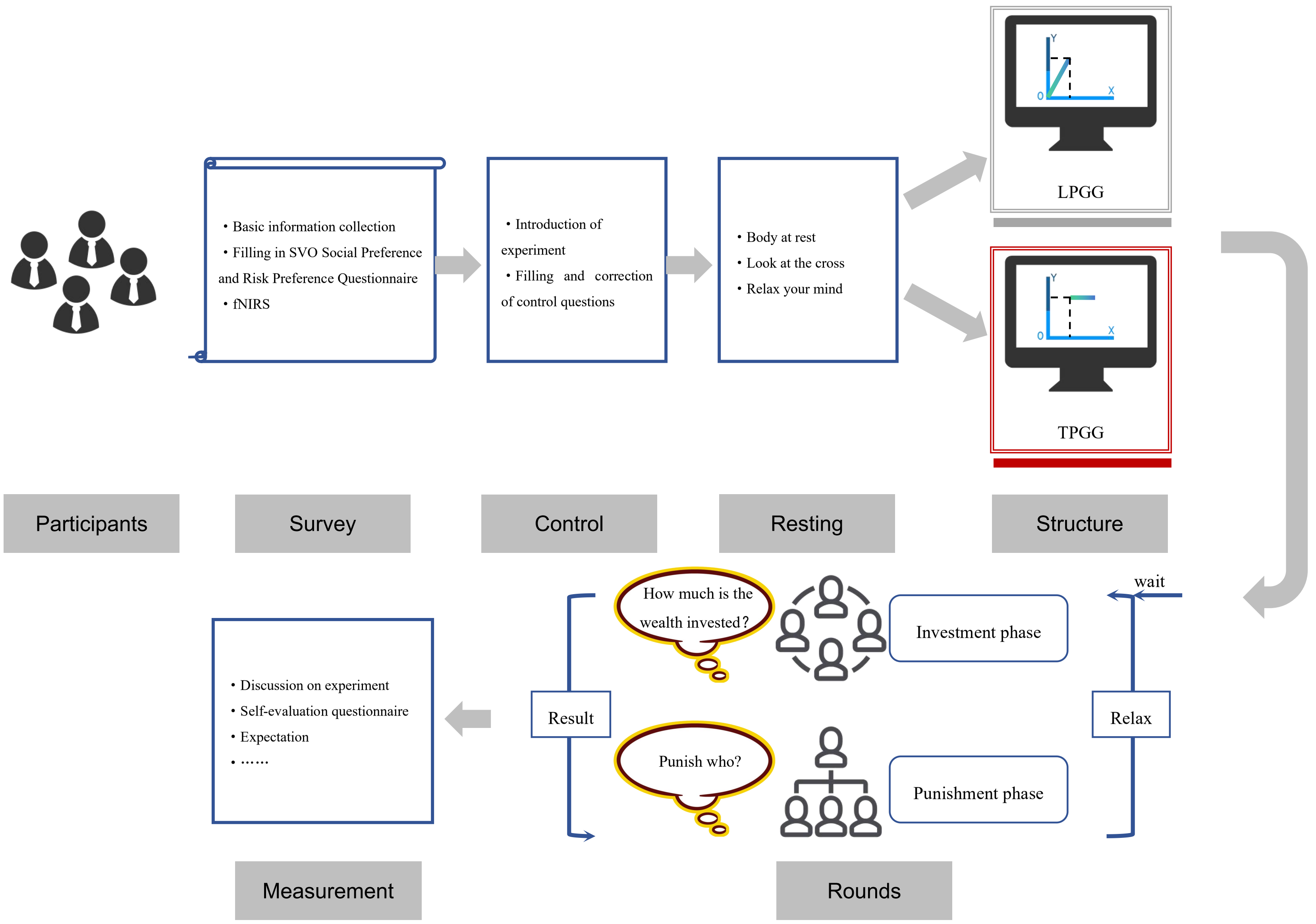
Basic information collection
Filling in SVO Social Preference and Risk Preference Questionnaire
fNIRS
Introduction of experiment
Filling and correction of control questions
Body at rest
Look at the cross
Relax your mind
LPGG
TPGG
Participants
Survey
Control
Resting
Structure
wait
How much is the wealth invested?
Investment phase
Punish who?
Punishment phase
Result
Relax
Discussion on experiment
Self-evaluation questionnaire
Expectation
……
Measurement
Rounds

**Supplementary Fig. 2**: **Experimental procedure.** Participants completed a pre-experiment questionnaire on the day before the main experiment, including demographic information, economic preference measures and social value orientation (SVO) [11, 12] (Section Pre-experiment questionnaire). On the experimental day, participants received task instructions and completed control questions to confirm their understanding of the game. They were then assigned their experimental condition and productivity type, followed by a 2-min resting-state baseline. The main task comprised four consecutive stages: preparation stage (rounds 1-6), first no punishment stage (NP1, rounds 7-21), punishment stage (P, rounds 22-36), and second no punishment stage (NP2, rounds 37-51). After completing the task, participants completed a post-experiment survey assessing their experience and feedback (Section **??**).

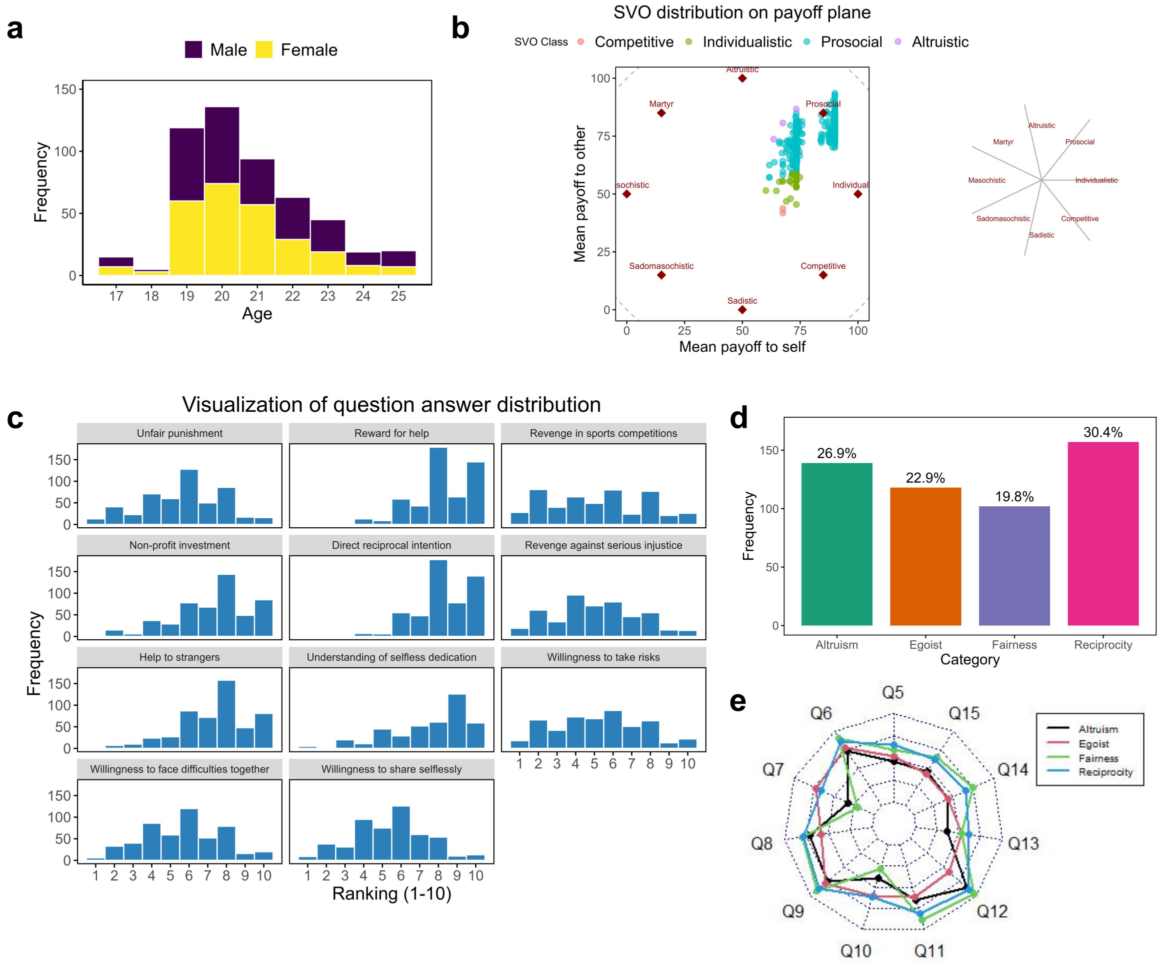

**Supplementary Fig. 3**: **Pre-experiment questionnaire results. a**, Age and gender distribution of participants. **b**, Distribution of social value orientation (SVO) on the self–other payoff plane, with most participants classified as prosocial. **c**, Response distributions across questionnaire items assessing punishment, helping, reciprocity, investment, risk preferences and related social preferences. **d**, Proportions of participants classified into altruism, egoism, fairness and reciprocity categories. **e**, Mean response profiles across questionnaire items for the four preference categories.

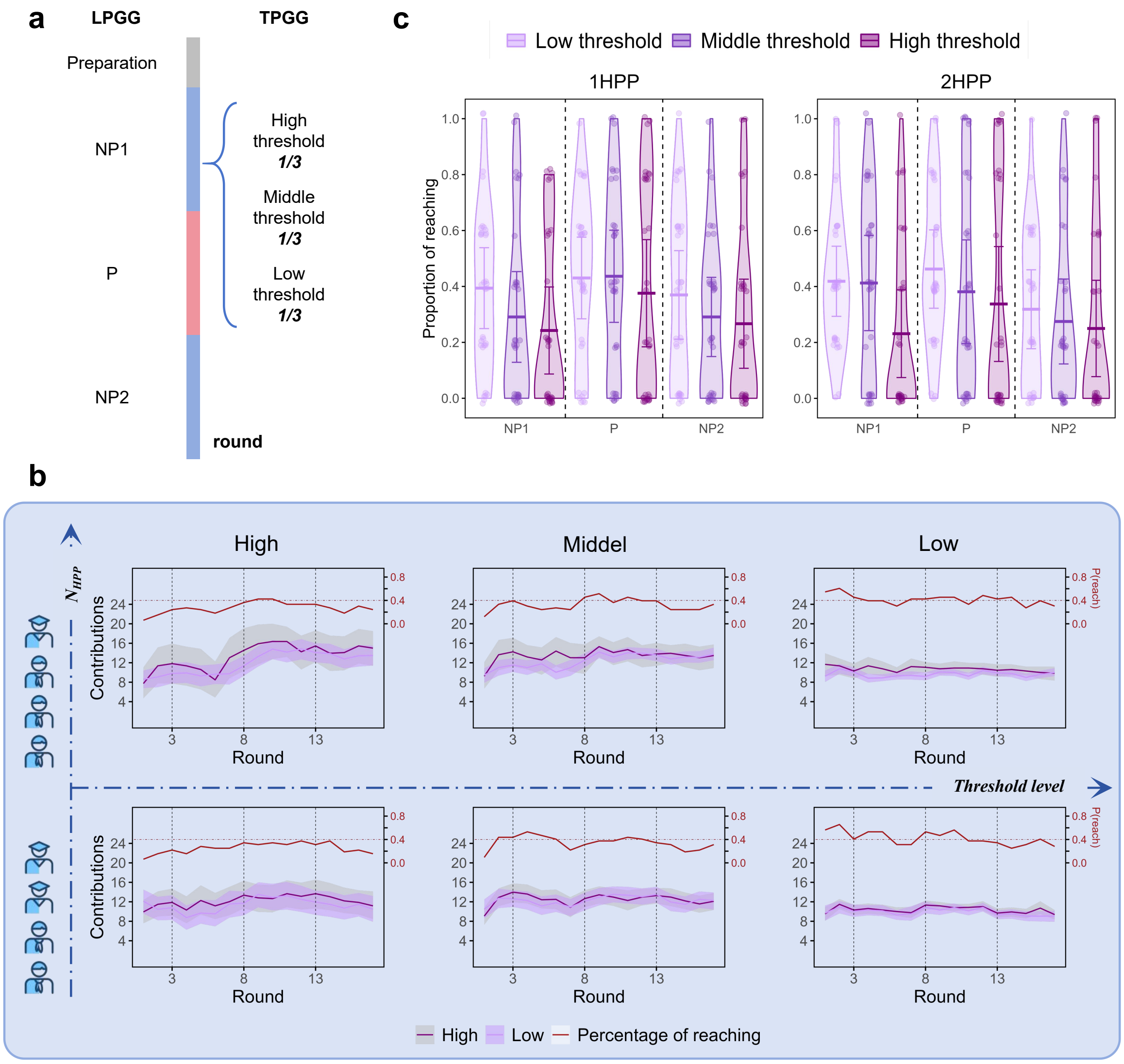

**Supplementary Fig. 4**: **Round structure and threshold-related outcomes in TPGG. a**, Experimental sequence across preparation, NP1, P and NP2 stages. In TPGG, high-, middle- and low-threshold rounds were equally represented within each stage. **b**, Round-by-round contributions and threshold-reaching rates for groups with one or two HPPs, shown separately for high, middle and low thresholds. **c**, Distribution of the proportion of rounds in which groups reached each threshold across NP1, P and NP2, separately for one- and two-HPP groups. Threshold attainment was generally higher during the P stage, particularly at the high and middle thresholds.

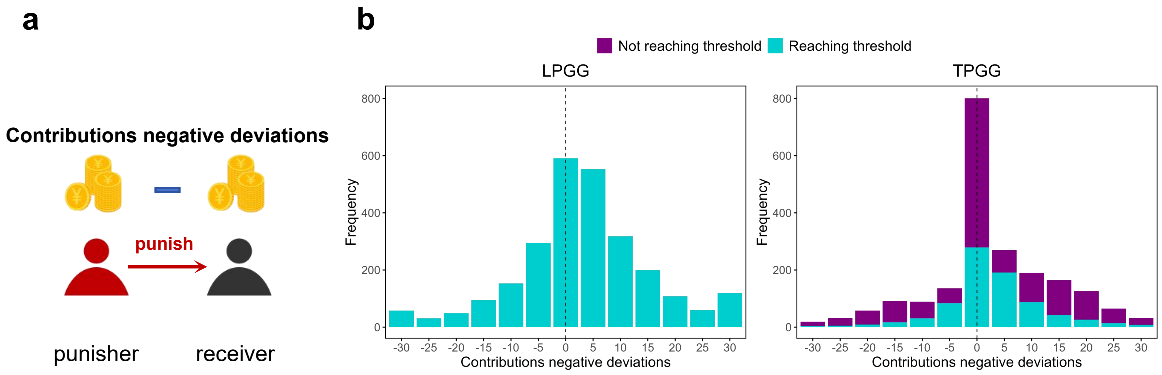

**Supplementary Fig. 5**: **Punishment frequence for a given deviation from the punisher's contribution. a**, Schematic definition of contribution deviation, calculated as the punisher's contribution minus the recipient's contribution within a given round. **b**, Distributions of contribution deviations for punishment events in LPGG and TPGG. In TPGG, punishment events are further separated according to whether the group reached the threshold. The mean contribution deviation was slightly smaller in TPGG than in LPGG (LPGG: $M = 3.137$, SD $= 12.145$; TPGG: $M = 2.578$, SD $= 11.244$). Within TPGG, the mean deviation was lower in rounds that failed to reach the threshold than in rounds that reached it (not reaching: $M = 2.494$, SD $= 12.499$; reaching: $M = 2.713$, SD $= 8.880$).

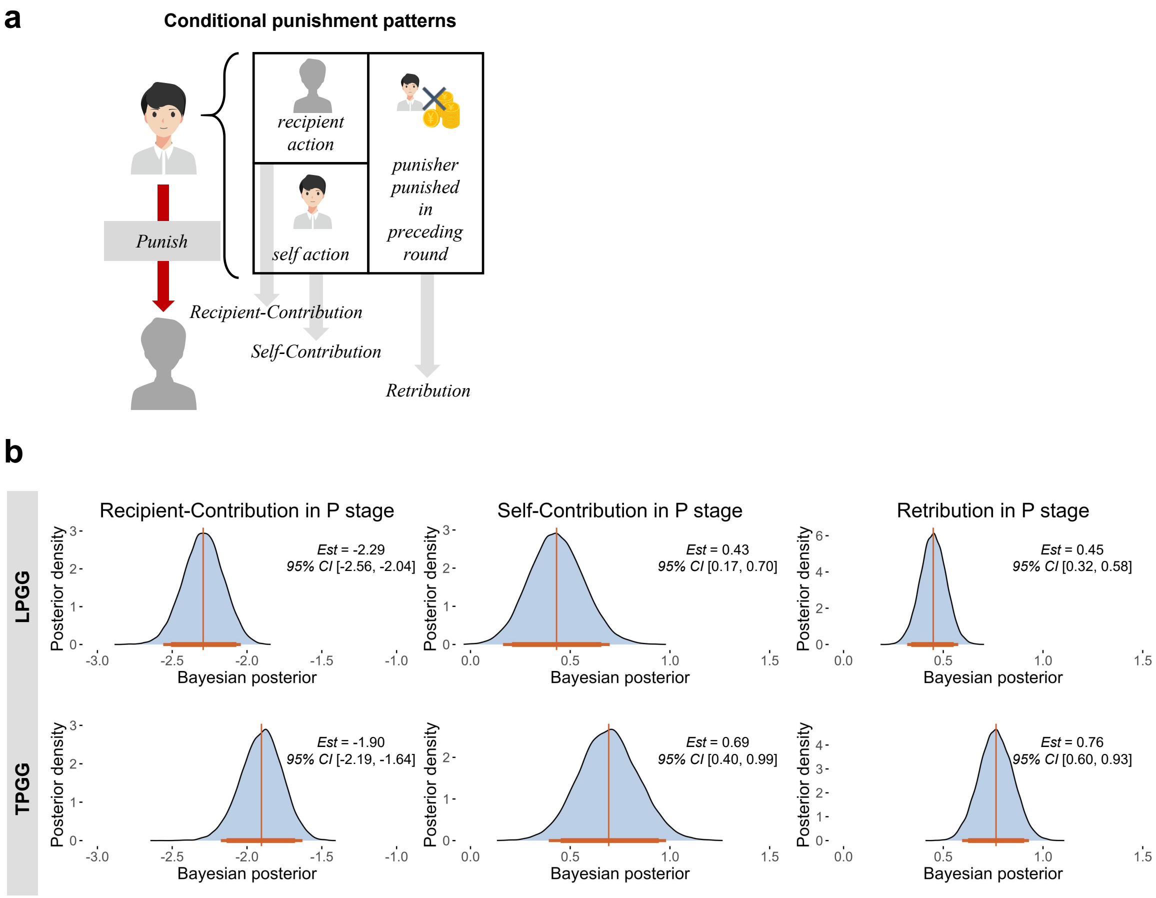

**Supplementary Fig. 6**: **Conditional punishment patterns across game structures. a**, Schematic of the conditional punishment analysis. Punishment decisions were modelled as a function of the recipient's contribution, the punisher's own contribution and the punishment received by the punisher in the preceding round (retribution). **b**, Posterior distributions of the corresponding regression coefficients during the P stage. In both games, lower recipient contributions were associated with greater punishments. TPGG showed a stronger negative association with recipient contribution and stronger positive associations with self-contribution and retribution than LPGG. Curves show posterior distributions, with posterior estimates and 95% credible intervals indicated.

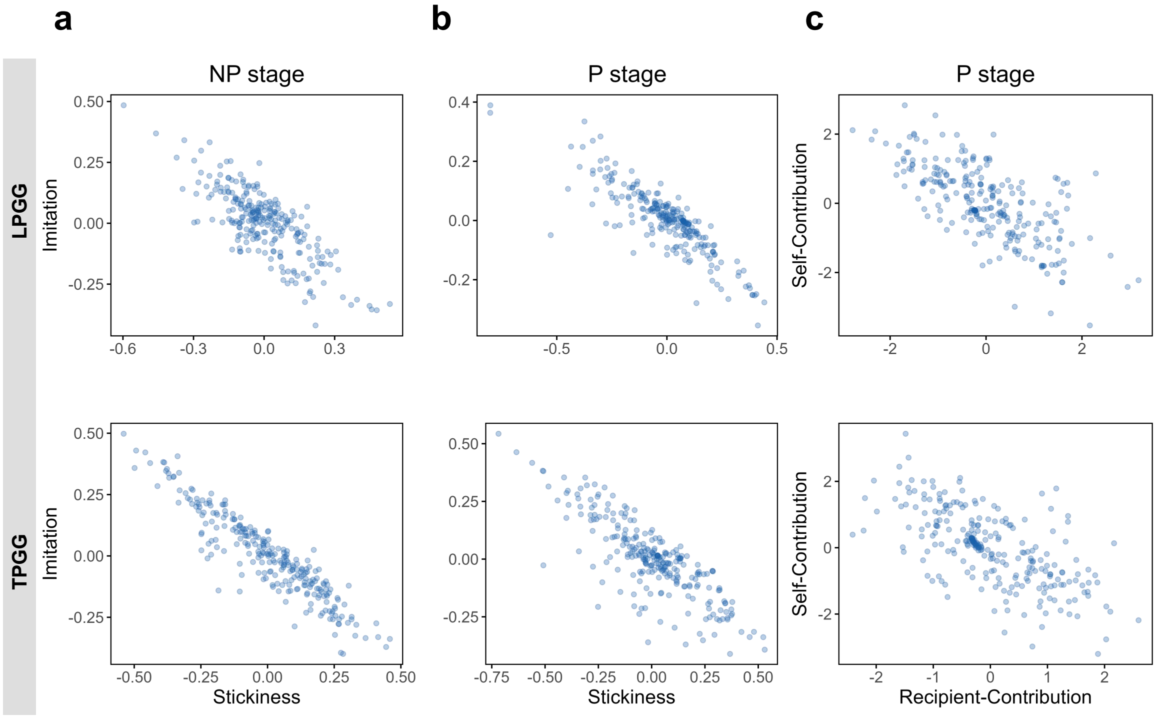

**Supplementary Fig. 7**: **Correlations among participant-specific random effects across game structures.** **a**, Relationship between random effects for stickiness and imitation during the NP stage. **b**, Relationship between random effects for stickiness and imitation during the P stage. **c**, Relationship between random effects for recipient-contribution and self-contribution effects during the P stage. In both LPGG and TPGG, each parameter pair showed a negative association.

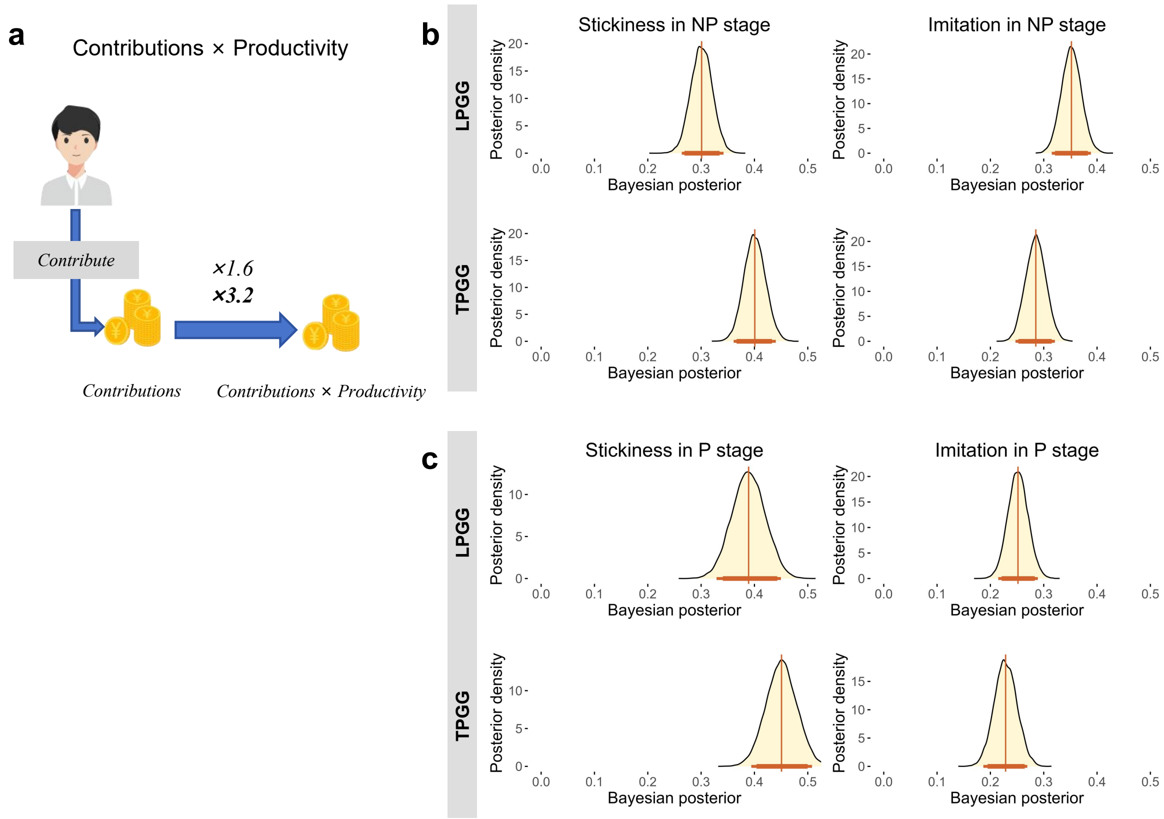

**Supplementary Fig. 8: Conditional contribution patterns based on productivity-weighted contributions.** **a**, Schematic of the productivity-adjusted contribution measure. Individual contributions were multiplied by the corresponding productivity to obtain productivity-weighted contributions, and the conditional contribution analysis was repeated using this measure. **b**, Posterior distributions of stickiness and imitation coefficients during the NP stage. **c**, Posterior distributions of the corresponding coefficients during the P stage. The results reproduced the main behavioral pattern: LPGG showed lower stickiness and higher imitation than TPGG, whereas punishment increased stickiness and reduced imitation in both game structures. Curves show posterior distributions, with posterior estimates and 95% credible intervals indicated.

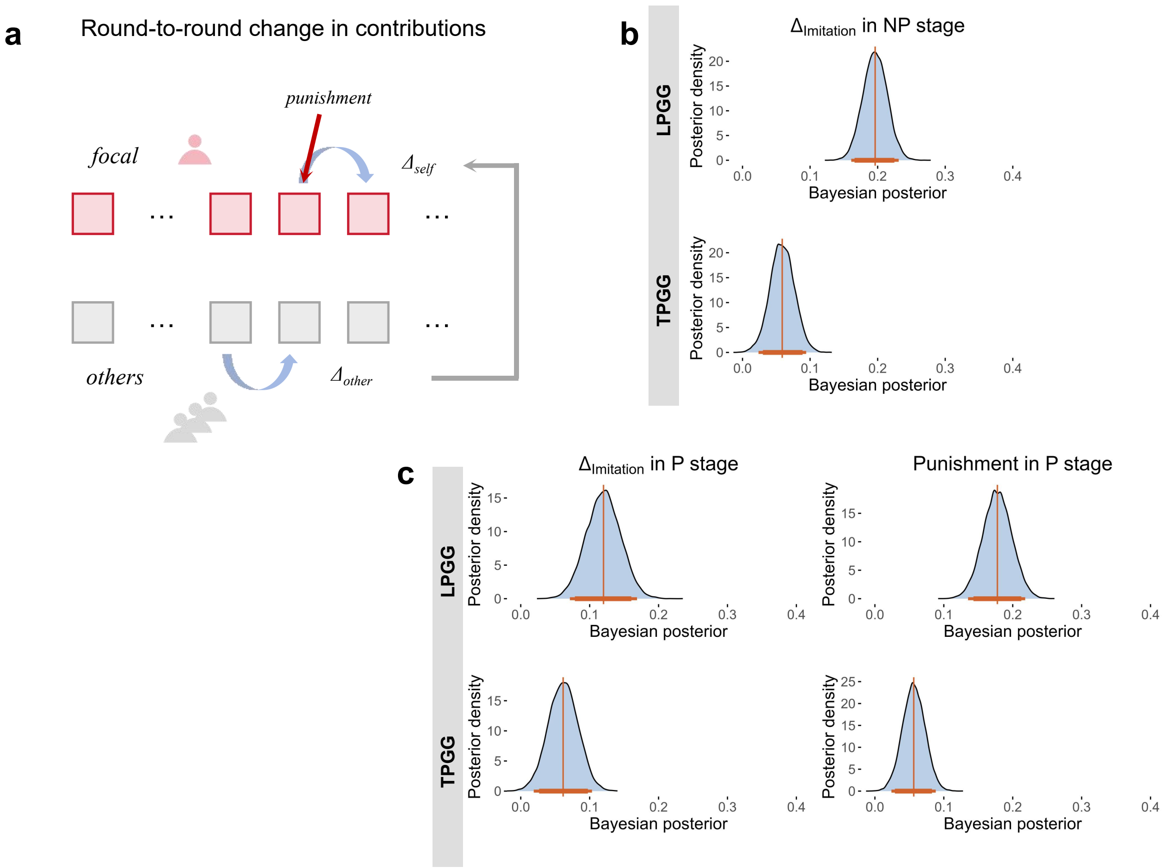

**Supplementary Fig. 9**: **Conditional contribution dynamics across game structures. a**, Schematic of the round-to-round contribution-change analysis. Contribution dynamics were defined as the change in contribution between consecutive rounds ($\Delta$) [13]. Changes in the focal participant's contribution were modelled as a function of changes in others' contributions; during the P stage, punishment received in the preceding round was additionally included as a predictor. **b**, Posterior distributions of the $\Delta_{\text{imitation}}$ coefficient during the NP stage. LPGG showed stronger dynamic responsiveness to changes in others' contributions than TPGG. **c**, Posterior distributions of $\Delta_{\text{imitation}}$ and punishment effects during the P stage. The $\Delta_{\text{imitation}}$ effect remained stronger in LPGG, and punishment exerted a larger positive effect on subsequent contribution changes in LPGG than in TPGG. Curves show posterior distributions, with posterior estimates and 95% credible intervals indicated.

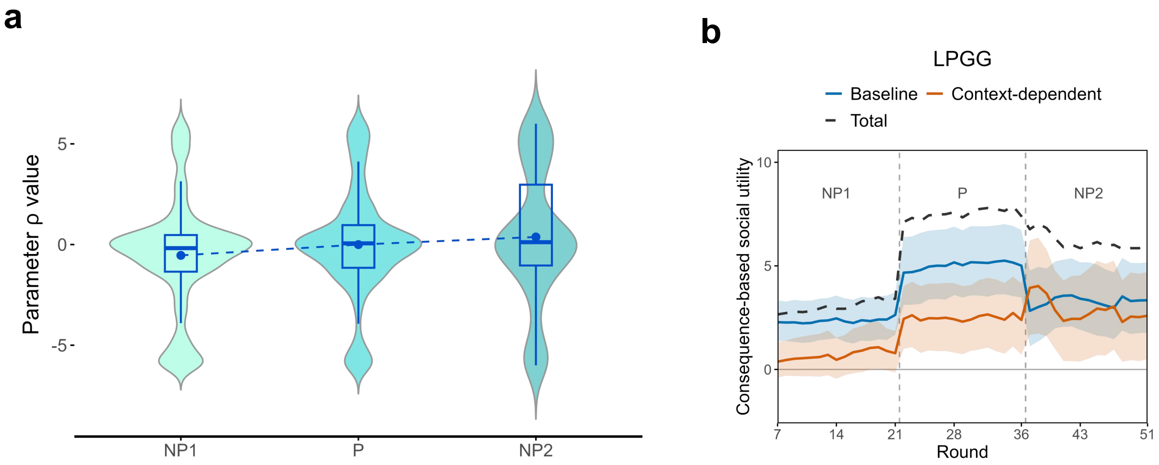

**Supplementary Fig. 10**: **Context-dependent modulation of consequence-based social value in LPGG. a**, Distribution of the context-dependent modulation $\rho$ across NP1, P and NP2. **b**, Round-by-round decomposition of consequence-based social utility in LPGG into the baseline component, the context-dependent component and their sum. Lines denote mean utility and shaded regions indicate 95% confidence intervals. The baseline component remained relatively stable across stages, whereas the context-dependent component increased after punishment became available and declined after its removal, producing corresponding changes in total consequence-based social utility.

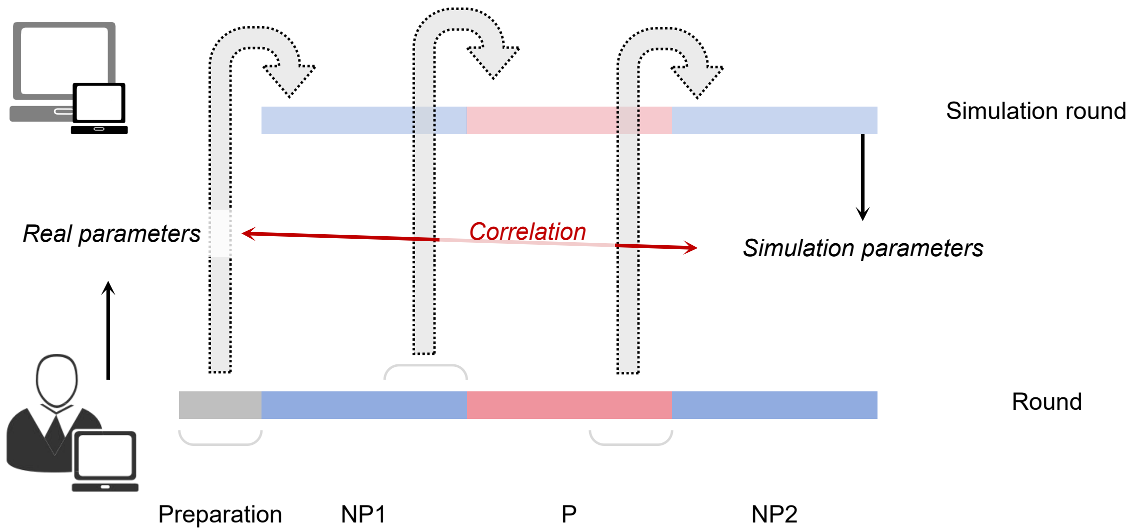

**Supplementary Fig. 11**: **Parameter-recovery procedure for the winning models.** Participant-specific parameters estimated from the observed data were used to generate synthetic contribution sequences for NP1, P and NP2 stages, with the social context initialized from the corresponding preceding experimental rounds. At each simulated round, contributions were sampled from the model-predicted choice distribution. The same winning model was then re-fitted to the simulated data to obtain recovered parameters. Parameter recovery was assessed by Pearson correlations between the generating and recovered parameter estimates.

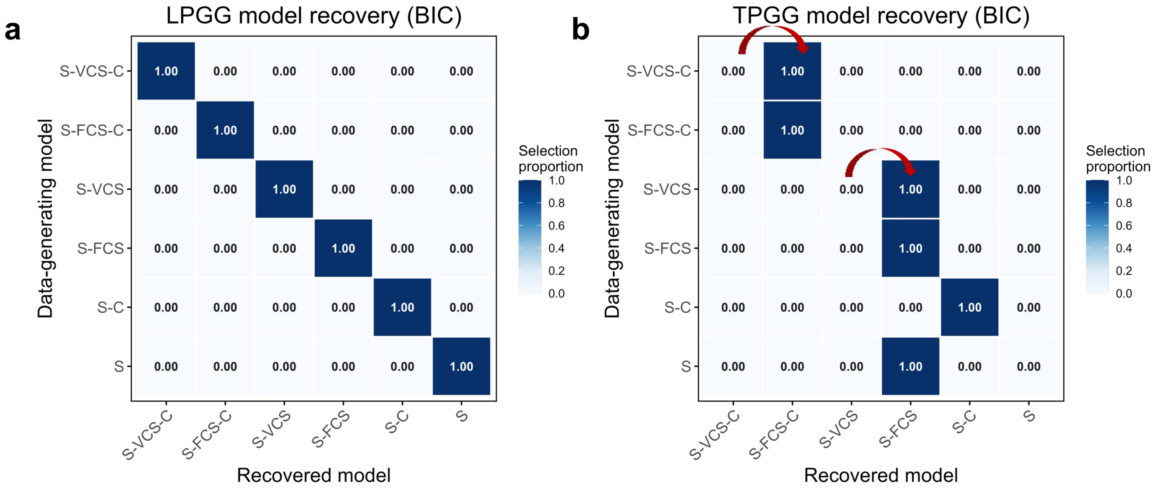

**Supplementary Fig. 12**: **Model-recovery analysis across game structures.** **a**, Confusion matrix for LPGG. All six candidate models were perfectly recovered from simulated datasets using BIC-based model selection. **b**, Confusion matrix for TPGG. Recovery was asymmetric: datasets generated by the $S$-$fCS$-$C$ and $S$-$fCS$ models were correctly identified, whereas datasets generated by the corresponding variable-weight models were consistently classified as their fixed-weight counterparts. Cell values indicate the proportion of simulated datasets assigned to each recovered model.

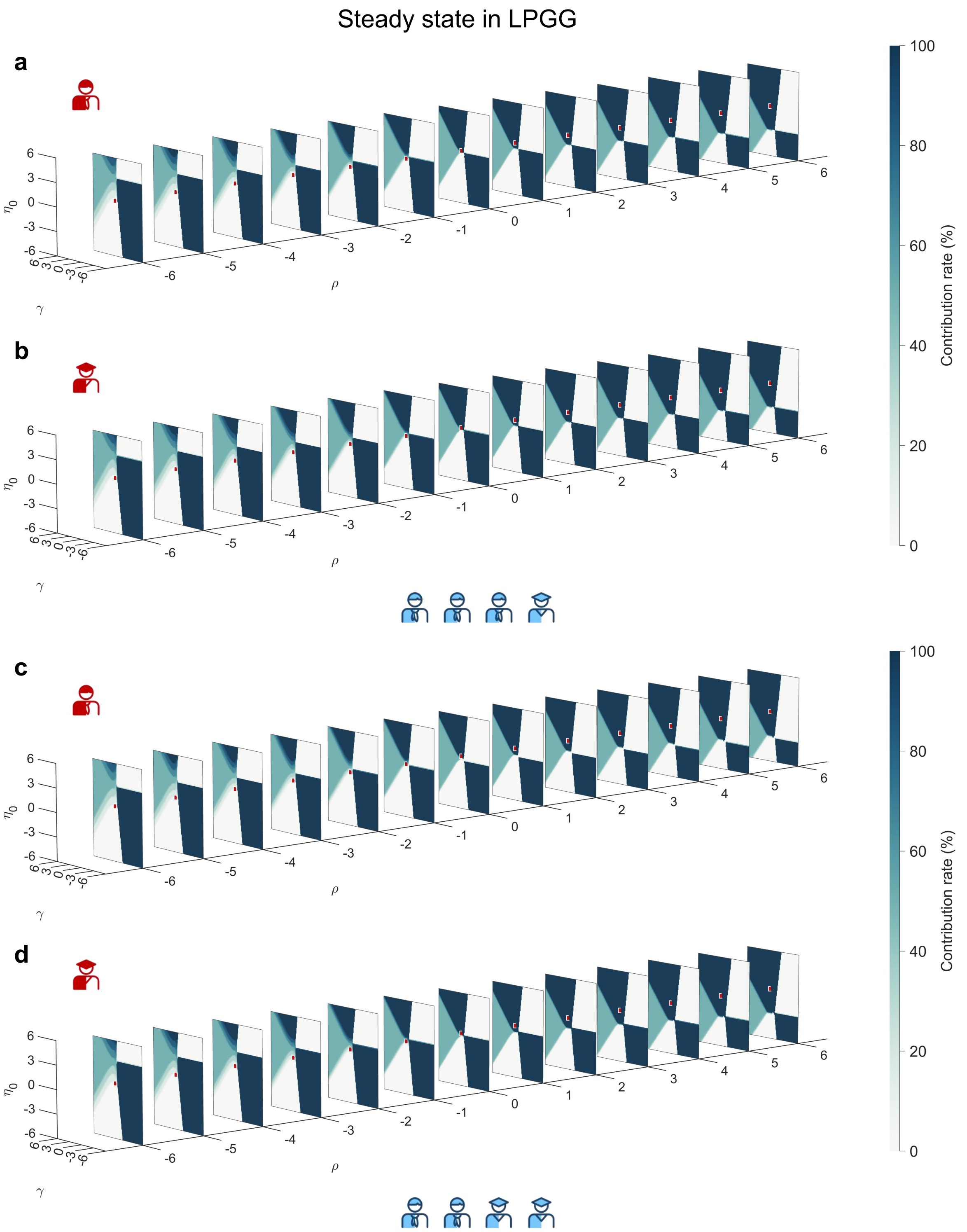

**Supplementary Fig. 13**: **Steady-state contribution rates across parameter regimes in LPGG. a–d**, Simulated steady-state contribution rates as a function of the baseline consequence-based social weight $\eta_0$, the context-dependent modulation $\rho$ and conformity weight $\gamma$. Results are shown separately for high- and low-productivity agents in groups containing either one high-productivity participant (**a,b**) or two high-productivity participants (**c,d**). Surface color denotes the steady-state contribution rate, and red markers indicate the parameters estimated from the observed data.

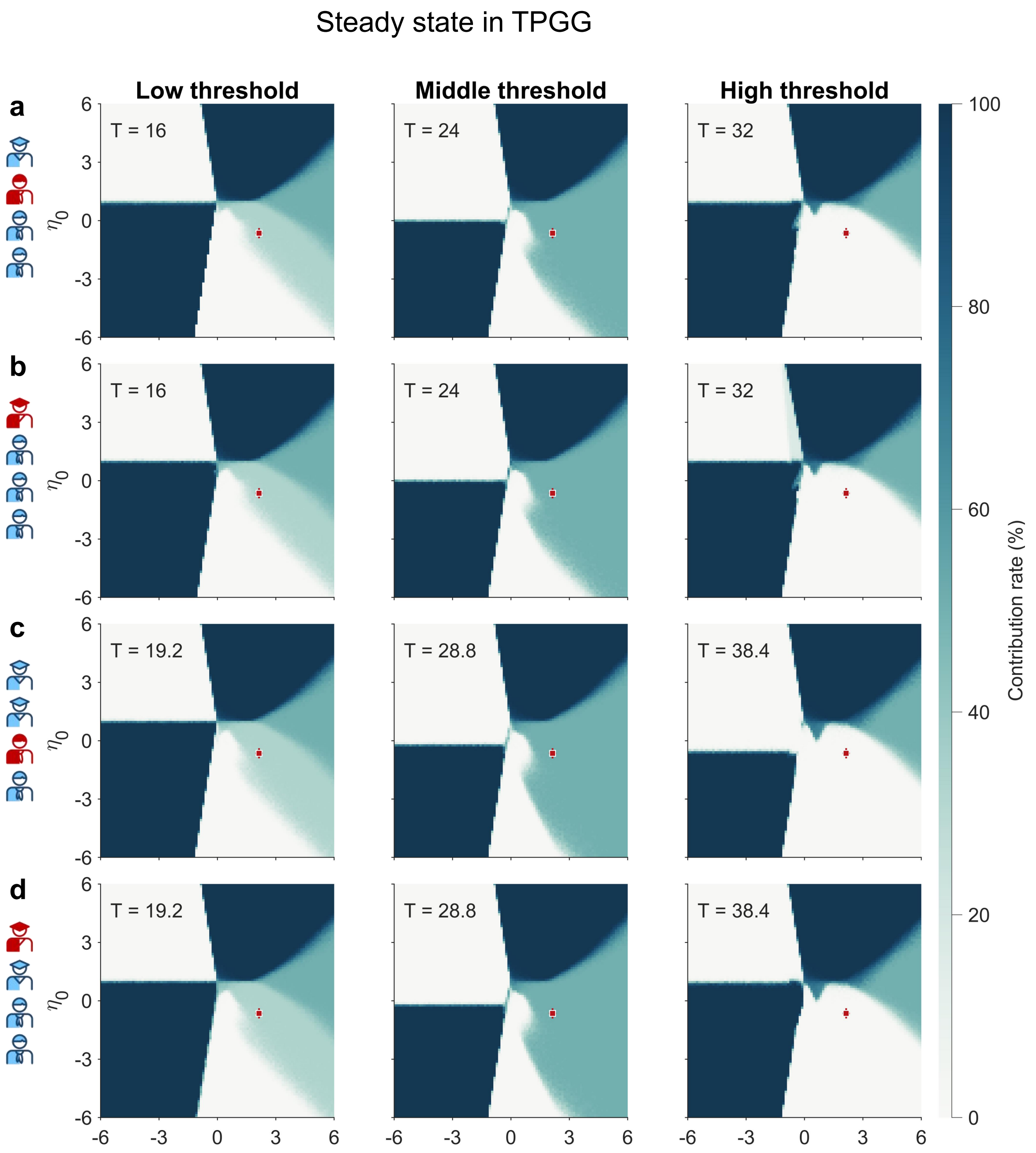

**Supplementary Fig. 14**: **Steady-state contribution rates across parameter regimes in TPGG. a–d**, Simulated steady-state contribution rates as a function of the baseline consequence-based social weight $\eta_0$ and conformity weight $\gamma$, shown separately for low-, middle- and high-threshold conditions. Results are shown separately for high- and low-productivity agents in groups containing either one high-productivity participant (**a,b**) or two high-productivity participants (**c,d**). Threshold $T$ were adjusted according to group productivity and threshold condition. Surface color denotes the steady-state contribution rate, and red markers indicate parameter estimates from the observed data.

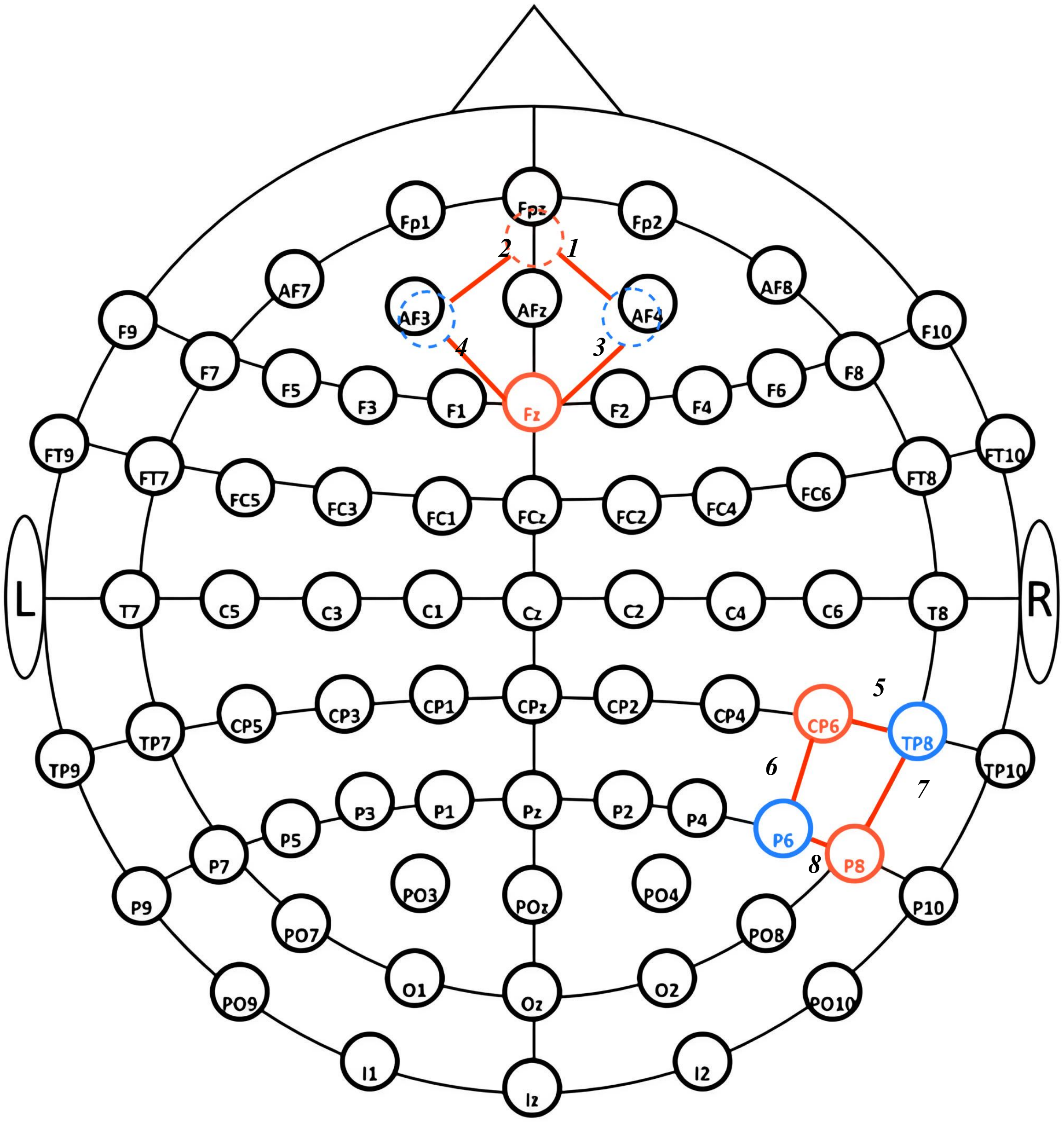

**Supplementary Fig. 15**: **fNIRS channel configuration.** Red and blue circles denote sources and detectors, respectively. Optodes were positioned according to the international 10–10 system to form eight channels covering the dorsomedial prefrontal cortex (dmPFC; channels 1–4) and right temporoparietal junction (rTPJ; channels 5–8). The posterior dmPFC source was aligned with Fz, and the source–detector separation was fixed at 3 cm.

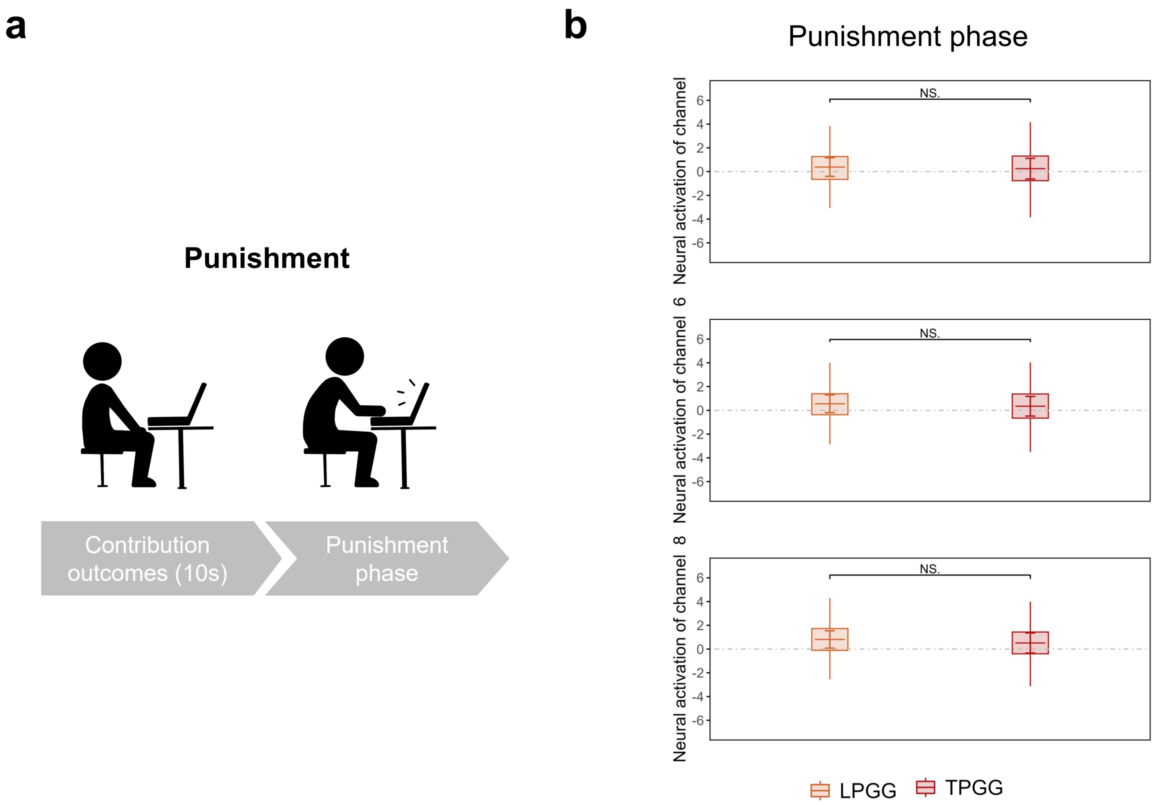

**Supplementary Fig. 16**: **Neural activation during the punishment phase.** **a**, Schematic of punishment-phase activation analysis. The 10-s contribution-outcome period preceding punishment served as the baseline, and HbO signals during the punishment phase were standardized using the corresponding baseline mean and standard deviation. **b**, Activation in rTPJ channels 5, 6 and 8 during punishment phases. Across NP1, P and NP2, no significant between-game differences were observed.

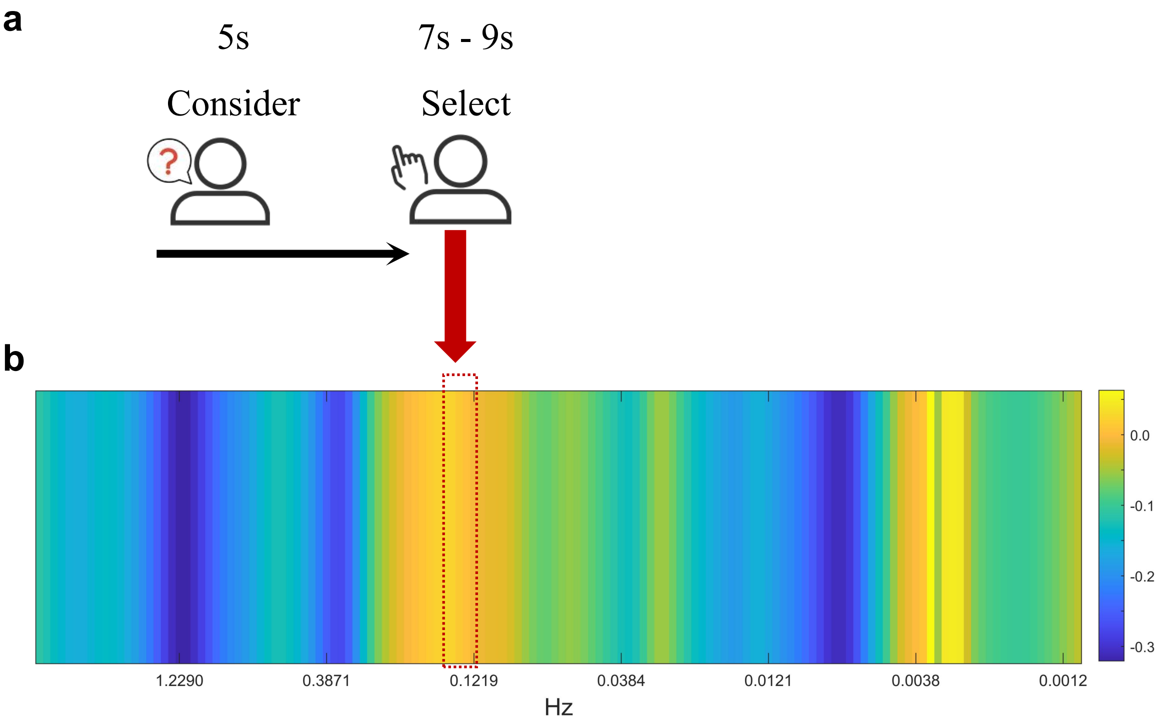

**Supplementary Fig. 17**: **Selection of the frequency of interest for INS analysis.** **a**, Schematic of the contribution phase, comprising a 5-s interval for considering followed by a 7–9-s interval for selecting. **b**, Mean task-related wavelet coherence across 142 frequency bins relative to the resting baseline. The frequency range showing the strongest task-related increase and overlapping the characteristic decision timescale was selected as the frequency of interest (0.1219–0.1536 Hz).

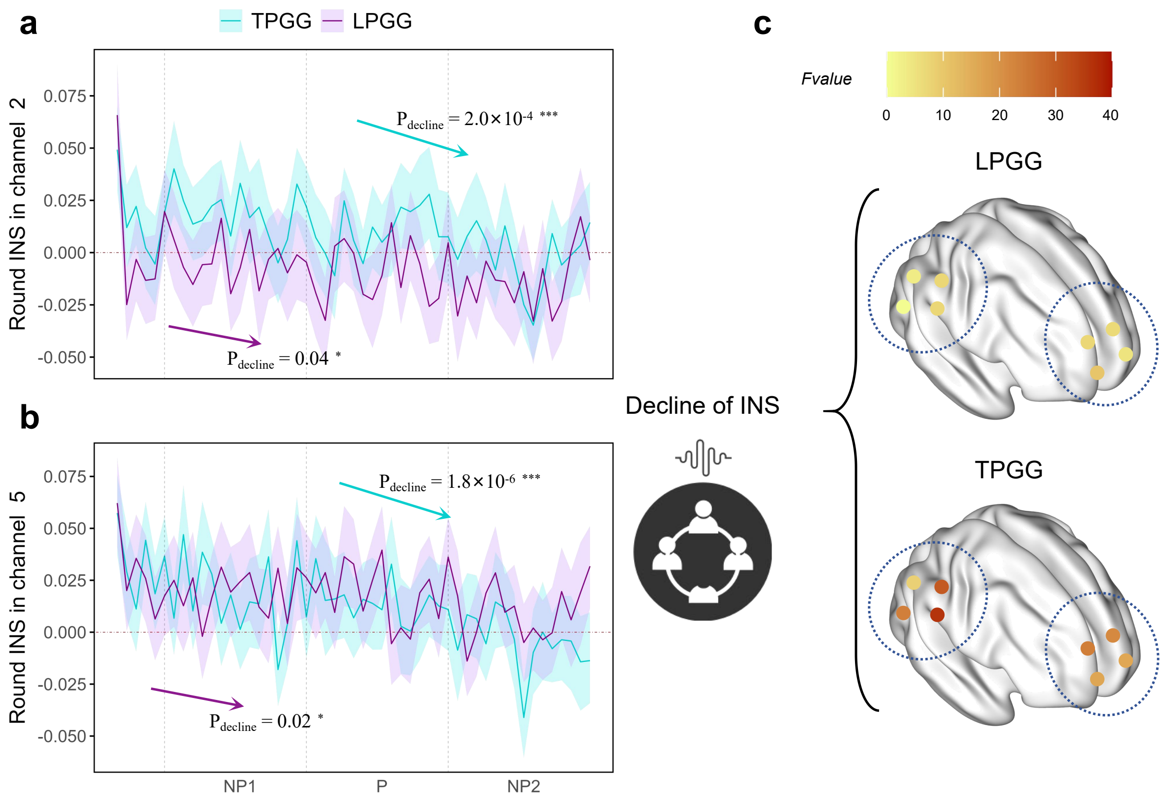

**Supplementary Fig. 18**: **Round-wise decline in group INS. a,b**, Changes in mean group-level INS across rounds for dmPFC channel 2 (**a**) and rTPJ channel 5 (**b**) in LPGG and TPGG. INS declined significantly over rounds in both game structures, with a steeper decrease in TPGG. Lines denote round-wise mean INS and shaded regions indicate uncertainty around the mean. **c**, Channel-wise $F$ statistics for the round-related decline in INS. The decline was generally stronger in TPGG than in LPGG across dmPFC and rTPJ channels.

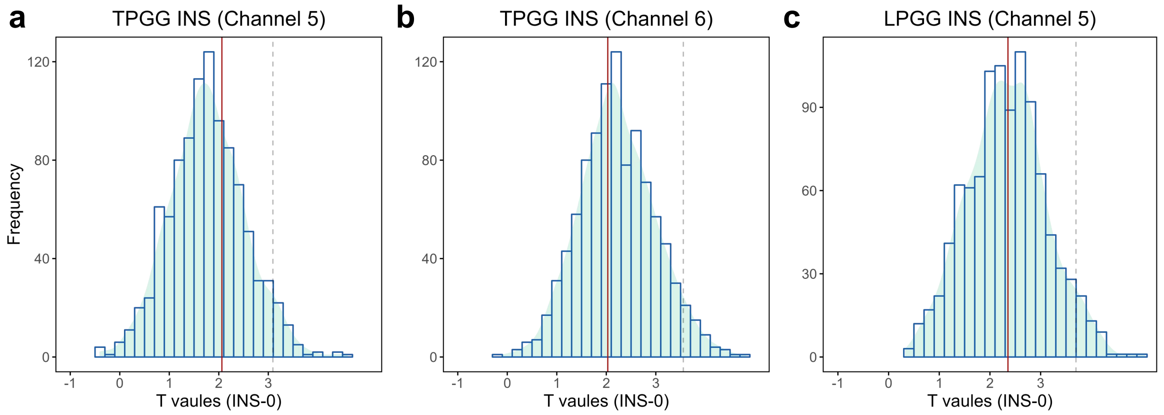

**Supplementary Fig. 19**: **Permutation tests of task-related INS. a,b**, Permutation distributions of rTPJ channels 5 (**a**) and 6 (**b**) synchrony in TPGG. **c**, Permutation distributions of rTPJ channels 5 in LPGG. Histograms show test statistics obtained from 1,000 pseudo-group permutations; red vertical lines indicate the statistics observed for the real interacting groups. Unlike dmPFC channel 2 in TPGG, the observed rTPJ effects did not exceed the permutation-based null distribution, indicating that these synchrony effects were not specific to real interacting groups.

### Supplementary Material 3

#### Pre-experiment questionnaire

1. Name [Fill-in-the-blank question] *

________________________

2. Date of Birth [Fill-in-the-blank question] *

________________________

3. Gender [Single-choice question] *

[ ]Male [ ]Female

The following statements describe what kind of person you are. Please indicate a number from 0 to 10——0 means [This statement does not describe me at all], and 10 means [This statement perfectly describes me]. You can also use any number between 0 and 10 to represent your position on this scale, such as 0, 1, 2, 3, 4, 5, 6, 7, 8, 9, 10.

4. Compared with others, I am a person who is willing to punish unfair behavior even if the cost of punishment is high. [Enter a number from 0 (does not match) to 10 (matches)] *

________________________

5. I make a special effort to help those who have helped me before. [Enter a number from 0 (does not match) to 10 (matches)] *

________________________

6. If someone is unfair to me in sports, I will be unfair to them in return. [Enter a number from 0 (does not match) to 10 (matches)] *

________________________

7. I am willing to spend time and money on non - profit things that seem meaningful to me, even if they do not directly benefit me. [Enter a number from 0 (does not match) to 10 (matches)] *

________________________

8. If someone does me a favor, I am willing to repay them. [Enter a number from 0 (does not match) to 10 (matches)] *

________________________

9. If I suffer a serious injustice, I will take revenge at all costs next time I have the chance. [Enter a number from 0 (does not match) to 10 (matches)] *

________________________

10. I am willing to help others, even if I think I will never meet them again. [Enter a number from 0 (does not match) to 10 (matches)] *

________________________

11. I don't understand why some people spend their whole lives fighting for a cause that doesn't directly benefit them. [Enter a number from 0 (does not match) to 10 (matches)] *

________________________

12. Compared with others, I am a person who is willing to take risks. [Enter a number from 0 (does not match) to 10 (matches)] *

________________________

13. When others are in trouble or an emergency, I am willing to face the difficulties with them without expecting any return. [Enter a number from 0 (does not match) to 10 (matches)] *

________________________

14. Compared with others, I am a person who usually likes to share with others without expecting something in return. [Enter a number from 0 (does not match) to 10 (matches)] *

________________________

The following are hypothetical scenario questions.

15. Imagine: I got lost while shopping in a strange city and asked a stranger for directions. The stranger offered to drive me to my destination. The drive took about 20 minutes, and the stranger spent about 100 yuan in total. The stranger didn't want money. I had six bottles of wine with me, with the following prices. I decided to give the stranger a bottle of wine as a thank - you gift. [Single-choice question] *

[ ]25 [ ]50 [ ]75 [ ]100 [ ]125 [ ]150

16. Imagine: I won 10,000 yuan in the lottery. Considering my current situation, I would donate ___________ yuan to a charity. [Fill-in-the-blank question] *

17. Imagine: I participated in an activity with a stranger and we won 100 yuan. The rule is that the stranger proposes a plan to divide the 100 yuan. After I know the plan, I decide whether to accept or reject it. If I accept, the money will be divided according to the plan; if I reject, neither of us will get anything. After thinking, I think the other party should offer me at least ___________ yuan for me to accept. [Fill-in-the-blank question] *

18. Imagine: I won a competition and can choose between two payment methods: (1) a lottery where I have a 50% chance of getting 1000 yuan and a 50% chance of getting nothing; (2) a fixed bonus between 0 and 1000 yuan. The fixed bonus should be at least ___________ yuan for me to prefer it to the lottery. [Fill-in-the-blank question] *

Imagine: I am doing an activity with a stranger. The rule is that we each start with 20 yuan. First, the other party can transfer any integer amount from 0 to 20 yuan to me, that is, 0, 1, 2, ..., up to 20. The amount transferred to me will be tripled and added to my account. Then I have the opportunity to transfer any integer amount back to the other party. The amount does not double in this process, and the maximum amount I can transfer depends on the total amount in my account after receiving the transfer from the other party.

19. Suppose the other party transfers 5 yuan to me. I have $20 + 3\times5 = 35$ yuan, and the other party has 20-5=15 yuan. I decide to transfer back ___________ yuan. [Fill-in-the-blank question] *

20. Suppose the other party transfers 10 yuan to me. I have $20 + 3\times10 = 50$ yuan, and the other party has $20 - 10 = 10$ yuan. I decide to transfer back ___________ yuan. [Fill-in-the-blank question] *

21. Suppose the other party transfers 15 yuan to me. I have $20 + 3\times15 = 65$ yuan, and the other party has $20 - 15 = 5$ yuan. I decide to transfer back ___________ yuan. [Fill-in-the-blank question] *

22. Suppose the other party transfers 20 yuan to me. I have $20 + 3\times20 = 80$ yuan, and the other party has $20 - 20 = 0$ yuan. I decide to transfer back ___________ yuan. [Fill-in-the-blank question] *

23. Suppose I were in the other person's party. First, I will transfer ___________ yuan. [Fill-in-the-blank question] *

Suppose you receive the following series of plans. For each plan, you need to select your favorite way of distribution to divide the wealth between you and another stranger.

#### Post-experiment survey

Dear students, please fill out the questionnaire carefully. If you have any questions, feel free to ask the experimenter in a timely manner.

1. Name [Fill-in-the-blank question] *

______________________

2. Your ID number [Single-choice question] *

[ ]A [ ]B [ ]C [ ]D

3. Your productivity [Single-choice question] *

[ ]1.6 [ ]3.2

4. How much personal wealth do you think a individual with a productivity of 3.2 should contribute per round? [Single-choice question] *

[ ]0 [ ]5 [ ]1 [ ]1 [ ]20 [ ]25 [ ]30

5. How much personal wealth do you think a individual with a productivity of 1.6 should contribute per round? [Single-choice question] *

[ ]0 [ ]5 [ ]1 [ ]15 [ ]20 [ ]25 [ ]30

6. You think [Single-choice question] *

[ ]Players with a productivity of 3.2 should contribute relatively more [ ]Players with a productivity of 1.6 should contribute relatively more [ ]The contributions should be the same

7. Your contribution strategy (You can rank multiple choices according to their importance) [Ranking question, please fill in the numbers in the brackets in order] *

[ ] I will contribute more if other participants contribute more, and contribute less if other participants contribute less

[ ] I will contribute more if other participants contribute less, and contribute less if other participants contribute more

[ ] Keep the contribution at a low level all the time, without changing with others

[ ] Keep the contribution at a high level all the time, without changing with others

[ ] Contribute randomly

[ ] Others

8. Given that contributing will cause losses, what is the reason for your contribution? [Fill-in-the-blank question] *

________________________

9. Which type of experiment do you prefer? [Single-choice question] *

[ ]Experiments with punishment [ ]Experiments without punishment

10. Briefly describe the reasons for your preference. [Fill-in-the-blank question] *

________________________

11. Your punishment strategy (You can rank multiple choices according to their importance) [Ranking question, please fill in the numbers in the brackets in order] *

[ ] Punish a participant if their contribution is lower than a certain value

[ ] Punish a participant if their contribution is higher than a certain value

[ ] Punish a participant if their contribution is not within a certain range

[ ] Punish a participant if their contribution is different from that of most people

[ ] Punish randomly

[ ] Other punishment methods

12. Given that punishing will cause losses, what is the reason for your punishment? [Fill-in-the-blank question] *

________________________

13. In the experiment, you are more inclined to [Multiple-choice question] *

[ ]Punish individuals with a productivity of 1.6

[ ]Not punish individuals with a productivity of 1.6

[ ]Punish individuals with a productivity of 3.2

[ ]Not punish individuals with a productivity of 3.2

[ ]Have no preference

14. In the experiment, did you try to estimate the contributed wealth of other participants? [Single-choice question] *

[ ]Yes [ ]No (Please skip to question 17)

15. How accurate do you think your estimate is? [Enter a number from 0 (inaccurate) to 6 (accurate)] *

______________________

16. What methods did you try to reach the threshold in the experiment? [Coordination mechanism] [Fill-in-the-blank question] *

______________________

17. What methods did you use to avoid being punished by others in the experiment? [Fill-in-the-blank question] *

______________________

#### Control questions

##### Control questions 1

**LPGG–2HPP**

To assist you in better understanding the experimental rules, please answer the following control questions. Input your calculated results (in wealth points)into the input box.

I. Investment Stage

The initial wealth in the investment stage is 30 for everyone. The payoff varies depending on how you allocate your 30 wealth points. Assume your productivity is 1.6, and the productivity of the other three subjects are 3.2, 3.2, and 1.6.

(1)Suppose neither you nor any other group members contribute to the company.

Question 1: What is your final wealth?

Question 2: What is the total final wealth of the other three subjects?

(2)Suppose you and the other three subjects each contribute 30 wealth points.

Question 3: What is your final wealth?

Question 4: What is the total final wealth of the other three subjects?

(3)Suppose the other three subjects each contribute 15 wealth points to the company.

Question 5: If you make no additional contribution, what is your final wealth?

Question 6: If you contribute 15 wealth points additionally, what is your final wealth?

Question 7: If you contribute 30 wealth points additionally, what is your final wealth?

(4)Suppose you contribute 15 wealth points to the company.

Question 8: If the other three subjects make no contribution, what is your final wealth?

Question 9: If the other three subjects each contribute 30 wealth points, what is your final wealth?

(5)Suppose you contribute 15 wealth points to the company.

Question 10: If the three subjects with productivity of 3.2, 3.2, and 1.6 contribute 0, 30, and 30 wealth points respectively, what is your final wealth?

Question 11: If the three subjects with productivity of 3.2, 3.2, and 1.6 contribute 30, 30, and 0 wealth points respectively, what is your final wealth?

**LPGG–1HPP**

To assist you in better understanding the experimental rules, please answer the following control questions. Input your calculated results (in wealth points)into the input box.

I. Investment Stage

The initial wealth in the investment stage is 30 for everyone. The payoff varies depending on how you allocate your 30 wealth points. Assume your productivity is 1.6, and the productivity of the other three subjects are 3.2, 1.6, and 1.6.

(1)Suppose neither you nor any other group members contribute to the company.

Question 1: What is your final wealth?

Question 2: What is the total final wealth of the other three subjects?

(2)Suppose you and the other three subjects each contribute 30 wealth points.

Question 3: What is your final wealth?

Question 4: What is the total final wealth of the other three subjects?

(3)Suppose the other three subjects each contribute 15 wealth points to the company.

Question 5: If you make no additional contribution, what is your final wealth?

Question 6: If you contribute 15 wealth points additionally, what is your final wealth?

Question 7: If you contribute 30 wealth points additionally, what is your final wealth?

(4)Suppose you contribute 15 wealth points to the company.

Question 8: If the other three subjects make no contribution, what is your final wealth?

Question 9: If the other three subjects each contribute 30 wealth points, what is your final wealth?

(5)Suppose you contribute 15 wealth points to the company. Question 10: If the three subjects with productivity of 3.2, 1.6, and 1.6 contribute 0, 30, and 30 wealth points respectively, what is your final wealth?

Question 11: If the three subjects with productivity of 3.2, 1.6, and 1.6 contribute 30, 0, and 30 wealth points respectively, what is your final wealth?

**TPGG–2HPP**

To assist you in better understanding the experimental rules, please answer the following control questions. Input your calculated results (in wealth points)into the input box.

I. Investment Stage

The initial wealth in the investment stage is 30 for everyone. The payoff varies depending on how you allocate your 30 wealth points. Assume your productivity is 1.6, and the productivity of the other three subjects are 3.2, 3.2, and 1.6. There are three types of thresholds: 96, 144, and 192.

(1)Suppose neither you nor any other group members contribute to the company.

Question 1: What is your final wealth?

Question 2: What is the total final wealth of the other three subjects?

(2)Suppose you and the other three subjects each contribute 30 wealth points.

Question 3: If the threshold is 96, what is your final wealth?

Question 4: If the threshold is 144, what is your final wealth?

Question 5: If the threshold is 192, what is your final wealth?

(3)If the threshold is 144, suppose the other three subjects each contribute 15 wealth points to the company. Question 6: If you make no additional contribution, what is your final wealth?

Question 7: If you contribute 15 wealth points additionally, what is your final wealth?

Question 8: If you contribute 30 wealth points additionally, what is your final wealth?

(4)If the threshold is 96, suppose you contribute 15 wealth points to the company.

Question 9: If the other three subjects make no contribution, what is your final wealth?

Question 10: If the other three subjects each contribute 30 wealth points, what is your final wealth?

(5)If the threshold is 192, suppose you contribute 15 wealth points to the company.

Question 11: If the three subjects with productivity of 3.2, 1.6, and 3.2 contribute 0, 30, and 30 wealth points respectively, what is your final wealth?

Question 12: If the three subjects with productivity of 3.2, 1.6, and 3.2 contribute 30, 0, and 30 wealth points respectively, what is your final wealth?

**TPGG–1HPP**

To assist you in better understanding the experimental rules, please answer the following control questions. Input your calculated results (in wealth points)into the input box.

I. Investment Stage

The initial wealth in the investment stage is 30 for everyone. The payoff varies depending on how you allocate your 30 wealth points. Assume your productivity is 1.6, and the productivity of the other three subjects are 3.2, 1.6, and 1.6. There are three types of thresholds: 80, 120, and 160.

(1)Suppose neither you nor any other group members contribute to the company.

Question 1: What is your final wealth?

Question 2: What is the total final wealth of the other three subjects?

(2)Suppose you and the other three subjects each contribute 30 wealth points.

Question 3: If the threshold is 80, what is your final wealth?

Question 4: If the threshold is 120, what is your final wealth?

Question 5: If the threshold is 160, what is your final wealth?

(3)If the threshold is 120, suppose the other three subjects each contribute 15 wealth points to the company. Question 6: If you make no additional contribution, what is your final wealth?

Question 7: If you contribute 15 wealth points additionally, what is your final wealth?

Question 8: If you contribute 30 wealth points additionally, what is your final wealth?

(4)If the threshold is 80, suppose you contribute 15 wealth points to the company.

Question 9: If the other three subjects make no contribution, what is your final wealth?

Question 10: If the other three subjects each contribute 30 wealth points, what is your final wealth?

(5)If the threshold is 160, suppose you contribute 15 wealth points to the company.

Question 11: If the three subjects with productivity of 3.2, 1.6, and 1.6 contribute 0, 30, and 30 wealth points respectively, what is your final wealth?

Question 12: If the three subjects with productivity of 3.2, 1.6, and 1.6 contribute 30, 0, and 30 wealth points respectively, what is your final wealth?

#### Control questions 2

##### LPGG–2HPP

To assist you in better understanding the experimental rules, please answer the following control questions. Input your calculated results (in wealth points)into the input box.

II. Investment Stage

The initial wealth in the investment stage is 30 for everyone. The payoff varies depending on how you allocate your 30 wealth points. Assume your productivity is 3.2, and the productivity of the other three subjects are 3.2, 1.6, and 1.6.

(1)Suppose the other three subjects each contribute 15 wealth points to the company.

Question 1: If you make no additional contribution, what is your final wealth?

Question 2: If you contribute 15 wealth points additionally, what is your final wealth?

Question 3: If you contribute 30 wealth points additionally, what is your final wealth?

(2)Suppose you contribute 15 wealth points to the company.

Question 4: If the other three subjects make no contribution, what is your final wealth?

Question 5: If the other three subjects each contribute 30 wealth points, what is your final wealth?

##### LPGG–1HPP

To assist you in better understanding the experimental rules, please answer the following control questions. Input your calculated results (in wealth points)into the input box.

II. Investment Stage

The initial wealth in the investment stage is 30 for everyone. The payoff varies depending on how you allocate your 30 wealth points. Assume your productivity is 3.2, and the productivity of the other three subjects are 1.6, 1.6, and 1.6.

(1)Suppose the other three subjects each contribute 15 wealth points to the company.

Question 1: If you make no additional contribution, what is your final wealth?

Question 2: If you contribute 15 wealth points additionally, what is your final wealth?

Question 3: If you contribute 30 wealth points additionally, what is your final wealth?

(2)Suppose you contribute 15 wealth points to the company.

Question 4: If the other three subjects make no contribution, what is your final wealth?

Question 5: If the other three subjects each contribute 30 wealth points, what is your final wealth?

**TPGG–2HPP**

To assist you in better understanding the experimental rules, please answer the following control questions. Input your calculated results (in wealth points)into the input box.

II. Investment Stage

The initial wealth in the investment stage is 30 for everyone. The payoff varies depending on how you allocate your 30 wealth points. Assume your productivity is 3.2, and the productivity of the other three subjects are 3.2, 1.6, and 1.6. There are three types of thresholds: 96, 144, and 192.

(1)If the threshold is 192, suppose the other three subjects each contribute 15 wealth points to the company. Question 1: If you make no additional contribution, what is your final wealth?

Question 2: If you contribute 15 wealth points additionally, what is your final wealth?

Question 3: If you contribute 30 wealth points additionally, what is your final wealth?

(2)If the threshold is 96, suppose you contribute 15 wealth points to the company.

Question 4: If the other three subjects make no contribution, what is your final wealth?

Question 5: If the other three subjects each contribute 30 wealth points, what is your final wealth?

**TPGG–1HPP**

To assist you in better understanding the experimental rules, please answer the following control questions. Input your calculated results (in wealth points)into the input box.

II. Investment Stage

The initial wealth in the investment stage is 30 for everyone. The payoff varies depending on how you allocate your 30 wealth points. Assume your productivity is 3.2, and the productivity of the other three subjects are 1.6, 1.6, and 1.6. There are three types of thresholds: 80, 120, and 160.

(1)If the threshold is 160, suppose the other three subjects each contribute 15 wealth points to the company. Question 1: If you make no additional contribution, what is your final wealth?

Question 2: If you contribute 15 wealth points additionally, what is your final wealth?

Question 3: If you contribute 30 wealth points additionally, what is your final wealth?

(2)If the threshold is 80, suppose you contribute 15 wealth points to the company.

Question 4: If the other three subjects make no contribution, what is your final wealth?

Question 5: If the other three subjects each contribute 30 wealth points, what is your final wealth?

#### Control questions 3

To assist you in better understanding the experimental rules, please answer the following control questions. Input your calculated results (in wealth points)into the input box.

III. Punishment Stage

You need to sacrifice 2 wealth points to punish one person. For each punishment a punished subject receives from another person, the punished subject's wealth decreases by 6 points. If the final wealth is negative, please fill in the negative number.

Question 1: If a subject punishes the other three people and is punished by the other three people simultaneously, what is his final wealth in the punishment stage?

Question 2: If a subject does not punish anyone but is punished by the other three people, what is his final wealth in the punishment stage?

Question 3: If you do not punish anyone and the other three people do not punish you either, what is your final wealth in the punishment stage?

Question 4: If you do not punish anyone and 2 out of the other three people punish you, what is your final wealth?

Question 5: If you punish 2 out of the other three people and 1 out of the other three people punishes you, what is your final wealth?
